# A Halo of Reduced Dinoflagellate Abundances In and Around Eelgrass Beds

**DOI:** 10.1101/712612

**Authors:** Emily Jacobs-Palmer, Ramón Gallego, Ana Ramón-Laca, Emily Kunselman, Kelly Cribari, Micah Horwith, Ryan P. Kelly

## Abstract

Seagrass beds provide a variety of ecosystem services, both within and outside the bounds of the habitat itself. Here we use environmental DNA (eDNA) amplicons to analyze a broad cross-section of taxa from ecological communities in and immediately surrounding eelgrass (*Zostera marina*). Sampling seawater along transects extending alongshore outward from eelgrass beds, we demonstrate that eDNA provides meter-scale resolution of communities in the field. We evaluate eDNA abundance indices for thirteen major phylogenetic groups of marine and estuarine taxa along these transects, finding highly local changes linked with proximity to *Z. marina* for a diverse group of dinoflagellates, and for no other group of taxa. Eelgrass habitat is consistently associated with dramatic reductions in dinoflagellate abundance both within the contiguous beds and for at least fifteen meters outside, relative to nearby sites without eelgrass. These results are consistent with the hypothesis that eelgrass-associated communities have allelopathic effects on dinoflagellates, and that these effects can extend in a halo beyond the bounds of the contiguous beds. Because many dinoflagellates are capable of forming Harmful Algal Blooms (HABs) toxic to humans and other animal species, the apparent salutary effect of eelgrass habitat on neighboring waters has important implications for public health as well as shellfish aquaculture and harvesting.

## 1 INTRODUCTION

Seagrass species are ecosystem engineers throughout the world’s coastal zones (22), generating and sustaining habitat for a multitude of associated taxa (8). Additionally, these marine macrophytes provide a wide variety of essential ecosystem services that directly benefit humans, such as temporary carbon sequestration (10), nursery habitat for human food species (18), and coastal protection through sediment accretion and stabilization (39; reviewed in 34). Even beyond the boundaries of the habitats themselves, organisms and ecosystems may benefit from spillover effects of eelgrass beds; one such benefit is a proposed reduced exposure to pathogens (e.g. 26).

Eelgrass (*Zostera marina*) is the dominant seagrass along temperate coasts of the Northern Hemisphere (48). Recent worldwide declines in this species and other seagrass taxa are alarming (37; but see 47), and have been met with local protection measures in some cases, such as designation of seagrass as a ‘Habitat Area of Particular Concern’ (see **?**), as well as a Puget Sound ‘Vital Sign’ indicator species (40) and the target of ‘no net loss’ policies (33). Frequently, a tradeoff between eelgrass conservation and aquaculture is presumed when such conservation efforts compete with shellfish seeding grounds (19). However, commercially important species such as oysters are in fact often proximally associated with *Z. marina* beds in the wild; they may thus depend on services provided by the habitat, and vice versa (for example, see 14).

To examine how the surrounding biological community changes in the vicinity of *Z. marina* habitats, we use environmental DNA (eDNA) from water samples to survey organisms on a series of alongshore transects in the estuarine waters of Washington State during the late spring and summer. By a large margin, we find that dinoflagellates are the group most affected by *Z. marina*; spatial proximity to eelgrass habitat is associated with a taxonomically widespread decrease in dinoflagellate abundance for meters outside the borders of the contiguous beds.

Several mechanisms could explain our observations. Among other possibilities, the apparent decrease in dinoflagellates in and around eelgrass may be due to (1) an ecological edge effect, a neutral function of a change in habitat over space (5), (2) a biophysical effect, in which slower rates of water movement within eelgrass beds tend to settle dinoflagellates out of the plankton, (3) a predation effect, in which consumers of dinoflagellates are more abundant within eelgrass habitat, or (4) an allelopathic effect, in which eelgrass or its associated species deter dinoflagellates (e.g. 21).

Given the observational nature of our study, we cannot definitively distinguish among these putative mechanisms. We do, however, consider our observations in their context, and find that our data tend to lend support to the allelopathy hypothesis. If substantiated by further experimental evidence, the idea that eelgrass communities can suppress HAB-producing taxa in a halo of influence around the beds could strengthen connections between seagrass habitat and human health, particularly in native communities with elevated rates of shellfish consumption (54), and in the context of an increase in the number of shellfish harvesting closures (51; 31) with expanding toxigenic dinoflagellate distributions in the region of study. Thus, our findings may be important for management of both seagrass and shellfish in the many regions of the world where the two coincide.

## 2 METHODS

### 2.1 Environmental DNA sample collection

Environmental DNA sequenced at a single genetic locus can provide an assay of community composition consisting of many taxa. The design of the particular PCR primers used largely determines the taxonomic composition, but it is not uncommon to sequence hundreds of taxa from dozens of phyla in a given sampling effort. Here, we targeted a ca. 313 bp fragment of COI using a primer set (28) known to amplify a broad range of marine taxa including diatoms, dinoflagellates, metazoans, fungi, and others; this primer set is broadly used in ecological applications (e.g. 27; 12).

To determine the biological community composition within *Z. marina* beds and the surrounding habitat from eDNA, we sampled seawater from five sites in Puget Sound: Port Gamble, Case Inlet, Nisqually Reach, Skokomish, and Willapa Bay (Figure 1). We surveyed each location at three timepoints during late spring and summer, in May, July, and August of 2017. Specifically, we collected a 1 liter bottle of seawater immediately under the water surface, within 2 feet of the eelgrass canopy in a total water depth of 1-3 feet. We collected samples from eelgrass (located at the approximate center of contiguous beds, 47-90 meters inside the edge), from each point in a transect extending alongshore at 1, 3, 6, 10, and 15m outside the edge of the contiguous beds, and from a final location over bare substrate (located 16-670m outside the edge of the contiguous beds). The bed edge was defined as the point at which shoot density fell below 3 shoots/m^2^; see Figure S1 for a transect schematic and Table S1 for precise transect locations by site. All transects ran alongshore, with samples for a given transect gathered at a uniform tidal height (−1 to -3 feet mean lower low water level). Due to local geography and conditions, it was not always possible to gather all transect samples during each sampling event; a comprehensive list of samples gathered is given in Table S1. We kept samples on ice until processing by filtering 500 mL from each sample under vacuum pressure through a cellulose acetate filter with 47-mm diameter and 0.45-*μ*m pore size and stored the filter at room temperature in Longmire’s buffer (42). The final dataset consisted of 84 water samples.

**Figure 1:**
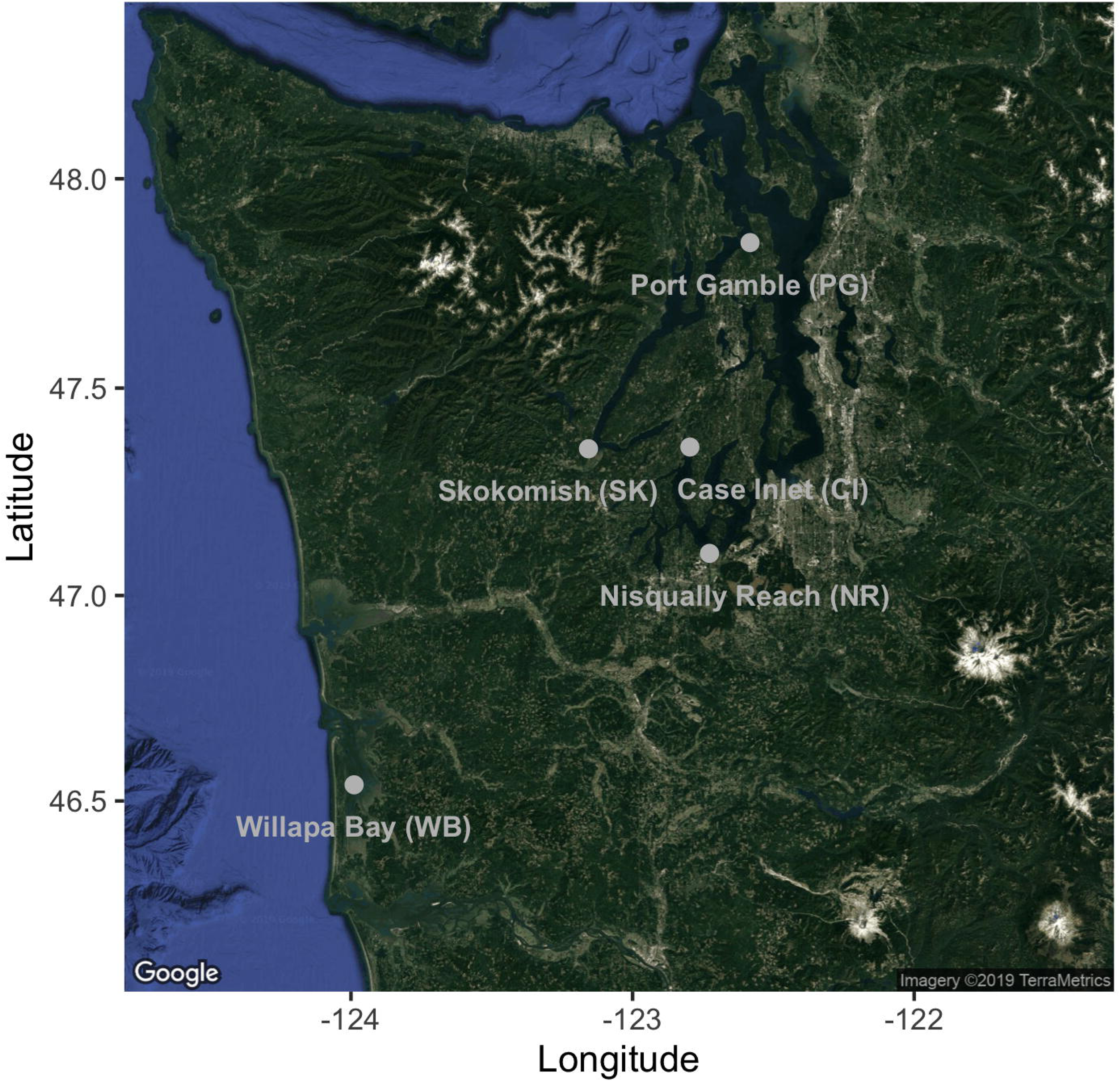
Nearshore sampling locations in Puget Sound and outer coast, Washington, USA. GPS coordinates are given in Supplemental Table 1.

### 2.2 Extraction and amplification

To extract DNA from the sample filters, we used a phenol:chloroform:isoamyl alcohol protocol (42). We incubated filter membranes at 65 °C for 30 min before adding 900 *μ*L of phenol:chloroform:isoamyl alcohol and shaking vigorously for 60 s. We conducted two consecutive chloroform washes by centrifuging at 14,000 rpm for 5 min, transferring the aqueous layer to 700 *μ*L chloroform, and shaking vigorously for 60 s. After a third centrifugation, 500 *μ*L of the aqueous layer was transferred to tubes containing 20 *μ*L 5 molar NaCl and 500 *μ*L 100% isopropanol, and frozen at -20 °C for approximately 15 h. Finally, all liquid was removed by centrifuging at 14,000 rpm for 10 min, pouring off or pipetting out any remaining liquid, and drying in a vacuum centrifuge at 45 °C for 15 min. We resuspended the eluate in 200 *μ*L water, and used 1 *μ*L of diluted DNA extract (between 1:10 and 1:400) as template for PCR.

To survey the eukaryotic organisms present in our samples, we ran and sequenced in triplicate PCR reactions from each of the 84 biological samples to distinguish technical from biological variance. To sequence many samples and their replicates in a single run while avoiding amplification bias due to index sequence, we followed a two-step PCR protocol (35). In the first step, we used a PCR reaction containing 1X HotStar Buffer, 2.5 mM MgCl2, 0.5 mM dNTP, 0.3 *μ*M of each primer, and 0.5 units of HotStar Taq (Qiagen Corp., Valencia, CA, USA) per 20 *μ*L reaction. The PCR protocol for this step consisted of 40 cycles, including an annealing touchdown from 62 °C to 46 °C (−1 °C per cycle), followed by 25 cycles at 46 °C. In the second step, following the 2-step PCR protocol given in O’Donnell et al. (35), we added 6 base-pair nucleotide tags to both ends of our amplicons prior to sequencing, allowing us to sequence multiple samples on the same MiSeq run. We allowed for no sequencing error in these tags; only sequences with identical tags on both the forward and reverse read-directions survived quality control. This gave us high confidence in assigning amplicons back to individual field samples.

Finally, we generated amplicons with the same replication scheme for both positive (kangaroo (genus *Macropus*) or ostrich (genus *Struthio*) tissue, selected because these species are absent from the sampling sites, and thus we could identify cross-contamination using reads from these taxa) and negative controls (molecular grade water), and verified by gel electrophoresis that negative controls contained no appreciable amount of DNA.

### 2.3 Sequencing

To prepare libraries of replicated, indexed samples and positive controls, we followed manufacturers’ protocols (KAPA Biosystems, Wilmington, MA, USA; NEXTflex DNA barcodes; BIOO Scientific, Austin, TX, USA). We then performed sequencing on an Illumina MiSeq (250-300 bp, paired-end) platform in four different sets of samples: two MiSeq V.2 runs and two MiSeq V.3 runs. We processed each batch separately through the initial bioinformatics analysis (see below). We employed hierarchical clustering on transects containing six PCR replicates sequenced across two different runs (three technical replicates per run derived from the same sampled bottle of water) and found that these samples were each others’ nearest neighbors (Figure S2); thus sequencing-run-level effects were negligible and we combined the data from the four sequencing runs.

### 2.4 Bioinformatics

We followed updated versions of previously published procedures for bioinformatics, quality control, and decontamination (23). This protocol uses a custom Unix-based script (11) calling third-party programs to perform initial quality control on sequence reads from all four runs combined, demultiplexing sequences to their sample of origin and clustering unique variants into amplicon sequence variants (ASVs) (30; 6). The output is a dataset including counts of each ASV per PCR replicate; ∼28M sequence reads from 19370 ASVs emerged from this step.

To address possible cross-sample contamination (see 45), we subtracted the maximum proportional representation of each ASV across all control samples (sequenced from extraction of kangaroo or ostrich tissue) from the respective ASV in field samples; 27 million reads from 19320 ASVs passed this step. After removing the two PCR replicates with an extremely low number of reads, we estimated the probability of ASV occurrence by performing site-occupancy modeling (44; 25). Following Lahoz-Monfort et al. 2016 and using the full Bayesian code for package rjags [REFREF CITE] provided by those authors, we modeled the probability of occupancy (i.e., true presence) for each of the unique sequence variants in our dataset. We treated replicate PCR reactions of each water bottle as independent trials, estimating the true-positive rate of detection (p11), false-positive rate (p10), and commonness (psi) in a Bayesian binomial model. We then used these parameters to estimate the overall likelihood of occupancy (true presence) for each ASV; those with low likelihoods (<20%) were deemed unlikely to be truly present in the dataset, and therefore culled. 25 million reads from 3143 ASVs survived this step.

Lastly, we removed samples whose PCR replicates were highly dissimilar: we calculated the Bray-Curtis dissimilarity amongst PCR replicates from the same bottle of water and discarded those with distance to the sample centroid outside a 95% confidence interval. Of 84 bottles of water collected, 3 technical replicates survived QC in 72 cases (86%), 2 replicates in 9 cases (11%), 1 replicate in 2 cases (2%), and zero replicates in a single case (1%) (Table S1). The final dataset of 24 million reads from 3142 ASVs comprised 83% of the original sequence reads.

All bioinformatic and analytical code is included in GitHub repositories [REFREF CITE], and provides the details of parameter settings in the bioinformatics pipelines used. Sequence and annotation information are included as well, and the former are deposited and publicly available in GenBank (upon acceptance; accession numbers will be provided in the published manuscript).

### 2.5 Taxonomy

To assign taxonomy to each ASV sequence, we followed the protocol detailed in Kelly et al. (23). Briefly, this protocol uses ‘blastn’ (7) on a local version of the full NCBI nucleotide database (current as of February 13, 2019), recovering up to 100 hits per query sequence with at least 85% similarity and maximum e-values of 10^−30^ (culling limit = 5), and reconciling conflicts among matches using the last common ancestor approach implemented in MEGAN 6.4 (20). Within MEGAN, we imposed an additional more stringent round of quality control to ensure sufficient similarity between query and database sequences by requiring a bit score of at least 450 (ca. 90% identical over the entire 313-bp fragment). Of the 24 million total reads in our dataset, we were able to annotate 4.1 million to the level of phylum or lower; the majority of the remaining reads had no BLAST hits meeting our criteria (7.6 million) or else did not receive taxonomic assignment due to insufficient similarity or conflicting BLAST hits (12.1 million). We use the annotated sequences in our taxonomic analyses below.

To examine patterns within the phylum Dinoflagellata, we further refined our annotations for these ASVs. Specifically, we considered the geographic range of taxa involved (restricting possible annotations to those taxa known from the North Pacific) and assigned taxonomy conservatively to the level of family (and genus, when possible) only in cases of >95% sequence identity between the subject and query sequence; ASVs we could not confidently assign to the level of family we excluded from further analyses. Multiple dinoflagellate sequences with identical amino-acid translations occurred within *Heterocapsa, Kareniaceae, Gymnodinium*, and *Hemotodinium*; to avoid pseudoreplication, we treated these as a single taxonomic unit (this choice did not affect the trends or significance of results).

### 2.6 Statistical Analysis

#### 2.6.1 Community Composition

To confirm the spatial resolution of our eDNA communities, we used non-metric multidimensional scaling (nMDS) ordination of eDNA indices for all ASVs within each technical replicate (38). To derive this index, we first normalized taxon-specific ASV counts into proportions within a technical replicate, and then transformed the proportion values such that the maximum across all samples is scaled to 1 for each taxon (24). Such indexing improves our ability to track trends in abundance of individual taxa in time and space by correcting for both differences in read depth among samples and differences in amplification efficiency among sequences; mathematically, it is equivalent to the Wisconsin double-standardization for community ecology as implemented in the vegan package for R (36). Using this index, we generated a single Bray-Curtis dissimilarity matrix for sequenced transect samples from each unique site/month combination and performed ordinations for each using the metaMDS function of vegan (36; 41) using a maximum of one-thousand random starts. We then created a single Bray-Curtis dissimilarity matrix for our entire dataset and apportioned variance by site, month, transect distance, and sample on the communities present using a PERMANOVA test (implemented with the adonis function in vegan (36)).

#### 2.6.2 Taxon-Habitat Associations

To examine the relative abundance of phyla in eelgrass habitat relative to bare substrate, we determined eDNA indices for the sum of sequences within each phylum at the two transect extremes (within-eelgrass vs. bare), calculating a relative eDNA abundance measure by subtracting the mean eDNA abundance index over bare substrate for each site-month combination from the corresponding mean eDNA abundance index in the eelgrass habitat. Positive values of this measure thus denote higher abundance in eelgrass, while negative values of this index indicate higher abundance over bare substrate. To assess the statistical significance of these phylum-level differences between habitat types, we compared the distributions of mean eDNA abundance indices for individual phyla in samples taken from eelgrass relative to their counterparts taken over bare substrate, using a paired Wilcoxon signed-rank test with Bonferroni correction for multiple comparisons.

#### 2.6.3 Dinoflagellate Distributions

To resolve the fine-scale patterns of dinoflagellates with respect to eelgrass, we focused on transects in which individual dinoflagellate taxa had overall high-abundance. To identify these transects, we took the grand mean of dinoflagellate taxon-specific eDNA indices for each technical replicate along transects at a given time and place, and used the k-means function of the R stats package (49) with k = 2 to separate transects from all present dinoflagellate taxa into two groups, high- and low-abundance, using unsupervised machine learning (Figure S3). Plotting data for individual taxa across transects for each site-month revealed episodic abundance of dinoflagellate sequences in time and space, as expected (Figure S4). A phylogeny built of dinoflagellate taxa from the high-abundance transects (Figure S5) confirmed that family- and genus-level taxonomic groups occupied monophyletic clades.

Eight transects identified by unsupervised clustering indicate high-abundance events. For these focal transects, we first compared the eelgrass habitat and bare substrate using a paired Wilcoxon signed-rank test of mean eDNA abundance index for each dinoflagellate taxon (here, having identified sequences to the level of family or genus, rather than grouping dinoflagellates together, as we have done above). Next, to determine whether dinoflagellate abundance measures at intermediate alongshore transect samples (1, 3, 6, 10, and 15 meters) were more closely associated with eelgrass habitat or bare substrate, we additionally performed Gaussian mixture modelling with two groups (46). We then used a Wilcoxon rank sum test to assess the significance of differences in the dinoflagellate eDNA abundance index distribution in the two groups produced by model-based clustering. To ensure that these groups did not result simply from spatial autocorrelation, we calculated Bray-Curtis dissimilarity based on eDNA abundance indices of all ASVs from adjacent points on each full transect. We tested the null hypothesis that spatial distance does not significantly influence Bray-Curtis dissimilarity using a Kruskall-Wallace test.

## 3 RESULTS

### 3.1 Community Composition

We assigned over 3,000 unique ASVs to thirteen eukaryotic phyla comprising a diverse set of single- and multicellular taxa including Arthropoda (arthropods), Annelida (annelid worms), Bacillariophyta (diatoms), Chlorophyta (green algae), Chordata (chordates), Cnidaria (cnidarians), Dinoflagellata (dinoflagellates), Echinodermata (echinoderms), Heterokonta (stramenopiles), Mollusca (molluscs), Nemertea (ribbon worms), Ocrophyta (brown algae), and Rhodophyta (red algae). This represents a broad – although by no means comprehensive – survey of eukaryotic communities in and around our sampled eelgrass beds.

Ordination via nMDS revealed consistent differentiation between eDNA communities across transects within a sampling site and date; technical replicates consistently clustered together. An example plot of samples gathered along the transect from eelgrass to bare substrate at Willapa Bay in July (Figure 2; all site/date plots shown in Figure S7) shows that the eelgrass community is quite dissimilar from other transect points along both axes. Moving away from eelgrass, most technical replicates of each sample bottle are fully distinguishable from those of other sample bottles (non-overlapping in ordination). For the instances in which complete transects were sampled at a given time and place (10) and all three technical replicates of a sample were available for analysis (60), 44 samples (73%) were similarly non-overlapping in ordination with all remaining transect points, demonstrating that despite proximity at the scale of meters, bottles of water contained eDNA evidence of distinct biological communities the majority of the time. Put differently, within-sample variance (reflecting laboratory-driven processes) was smaller than between-sample variance (reflecting biological as well as laboratory processes), hence providing resolution of communities at the scale of meters.

PERMANOVA apportioned the variance in Bray-Curtis distance among samples as follows: site (R2 = 0.18593, p = 0.001), month (R2 = 0.07909, p = 0.001), and transect distance (R2 = 0.02625, p = 0.001) each explain a significant portion of the variance in the dataset. Thus, despite strong effects of location and time, these results confirm that we can consistently distinguish nearshore eDNA communities (as sampled by our primers) at spatial scales of meters for each site and month of sampling. Moreover, we see a highly significant effect of proximity to eelgrass on the complement of organisms present.

**Figure 2:**
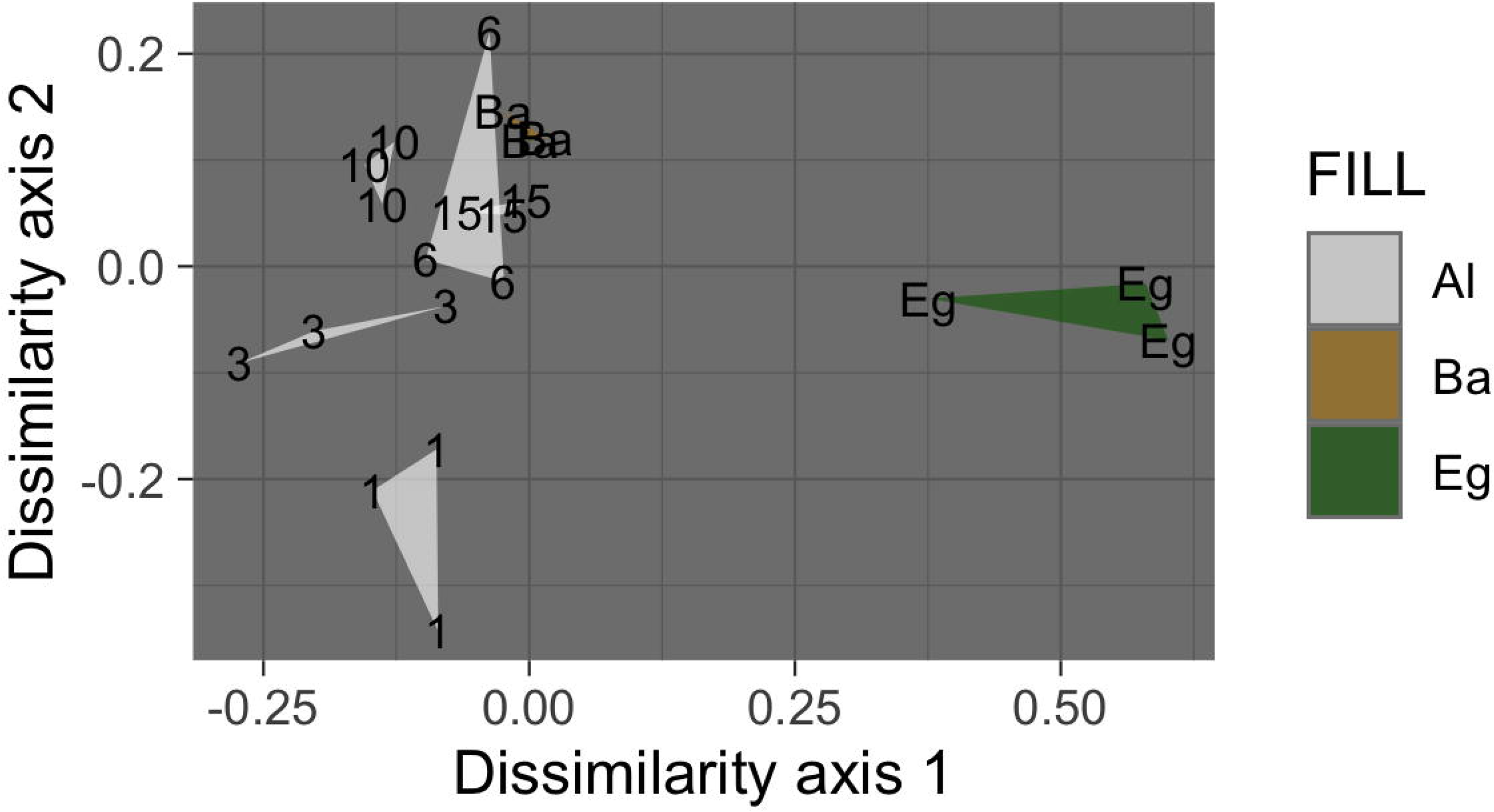
Example ordination plot of samples along a single transect from bare to eelgrass positions at Willapa Bay in July, 2017. Technical replicates of each biological sample are grouped as triangles. Alongshore transect samples are shown in white and labeled with distance from the contiguous eelgrass bed in meters (Al); the single sample taken above eelgrass (located 47-90 meters inside the edge of the contiguous beds) is shown in green (Eg), and that taken above bare substrate (located 16-670m outside the edge of the contiguous beds) is shown in brown (Ba).

### 3.2 Taxon-Habitat Associations

To determine the habitat preference of major taxa in our dataset at a course spatial scale, we classified ASVs to the level of phylum and plotted an index of their relative sequence abundance in eelgrass versus bare positions (Figure 3). Positive indices denote greater abundance in eelgrass, and negative indices in bare substrate. Across all sites and months, only dinoflagellates show a consistent and strong bias towards one habitat or another; they are nearly universally more abundant over bare substrate. Indeed, the negative association of dinoflagellates with eelgrass beds is the only significant change in phylum-specific abundance between the two habitat extremes after Bonferroni correction for multiple comparisons (p = 0.004; paired Wilcoxon signed-rank test). Other single-celled microalgae such as diatoms (Bacillariophyta) and green algae (Chlorophyta) have no significant relationship with eelgrass.

**Figure 3:**
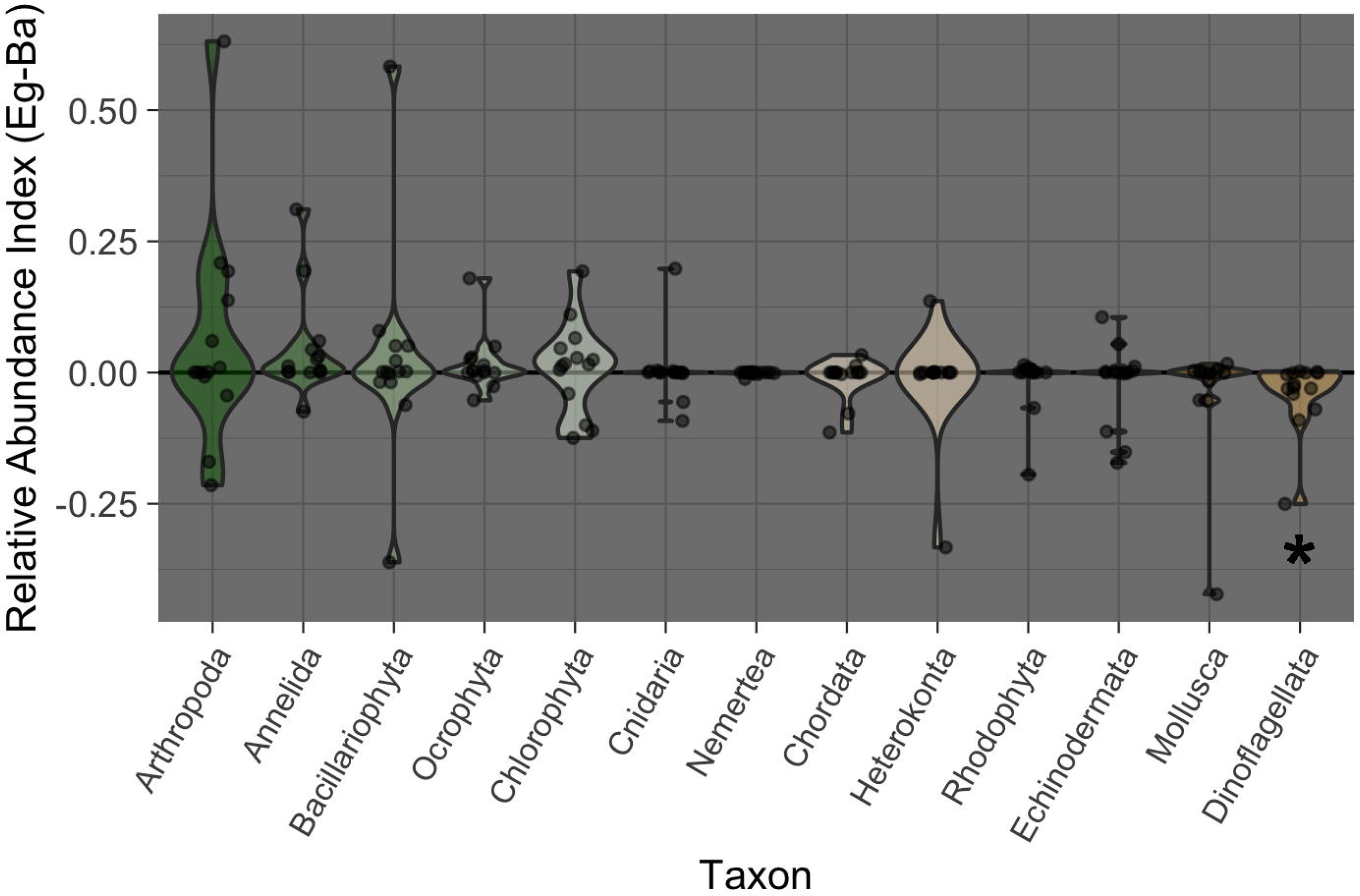
Habitat associations of sequences assigned to each phylum. Phyla are ordered and colored by mean relative eDNA abundance index (Eg-Ba: eDNA abundance index in eelgrass - eDNA abundance index over bare substrate). Greener samples on the left exhibit greater relative abundance in eelgrass, and browner samples on the right exhibit greater relative abundance on bare substrate. The central zero-line indicates no bias in abundance between habitat types. The violins are scaled to a fixed maximum width, and display the density of points within the interquartile range (shown by the violin area) around the median (central horizontal line). The significance of dinoflagellate habitat associations is denoted with an asterisk (paired Wilcoxon signed-rank test; p = 0.004).

~~~
## # A tibble: 2 × 2
## # Groups:        variable [2]
##  variable        p.value
## <chr>              <dbl>
## 1 Annelida       0.0310
## 2 Dinoflagellata 0.00403
~~~

### 3.3 Dinoflagellate Distributions

It is when dinoflagellates are plentiful relative to background levels that we have the power to identify trends in the abundance of individual taxa with respect to eelgrass habitat. To restrict our analysis to such periods, we used unsupervised machine learning (k-means clustering) to define a set of high- and low-abundance transects for each dinoflagellate sequence across all sites and months (Figure S3; between group sum of squares / total sum of squares = 67.7 %); eight transects from eight dinoflagellate taxa appeared in the high-abundance group. Their distributions are indeed highly local and episodic at the scale of our sampling, as expected (Figure S4).

In this subset of high-abundance transects, the negative interaction of eelgrass and dinoflagellates is taxonomically universal. The taxa represented include two unique variants from the genus *Heterocapsa*, single variants from the genera *Karlodinium, Alexandrium, Protoceratium*, and *Gymnodinium*, and two Kareniaceae family variants of unknown genus, all of which contain known or suspected HAB species (IOC Harmful Algal Bloom Programme and the World Register of Marine Species). All are also heavily biased towards bare substrate, relative to eelgrass (Figure 4; Wilcoxon signed-rank test, p < 0.002).

**Figure 4:**
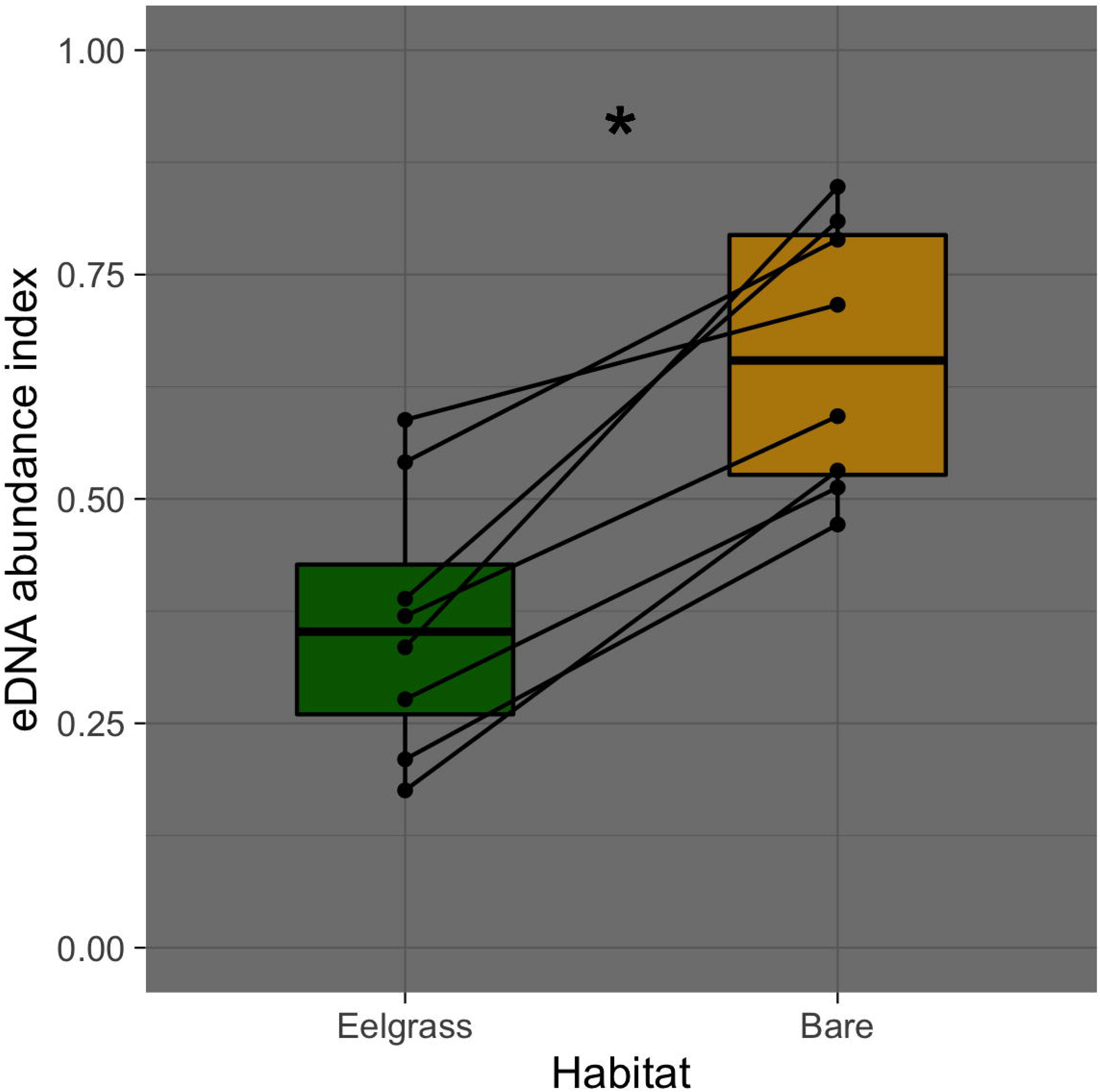
Habitat preferences of dinoflagellate sequences at site-months in which each taxon occurs at high abundance. eDNA abundance indices from Eelgrass samples are shown in green, and those from bare samples are shown in brown. The significance of paired differences in eDNA abundance indices for these transect extremes is denoted with an asterisk (Wilcoxon signed-rank test; p < 0.002).

After demonstrating a preference of all dinoflagellate taxa towards the bare substrate extreme (when highly-abundant), we then characterized their patterns as a function of distance from the edge of the contiguous eelgrass beds, using data from entire transects (Figure 5). Examining all points alongshore – and hence, controlling for substrate depth – we found that dinoflagellate eDNA abundance indices at the 1, 3, 6, 10, and 15m positions grouped with those at the eelgrass position in model-based clustering. Additionally, the eDNA abundance index of all high-abundance dinoflagellate taxa at these six transect points together differed significantly from bare substrate (Wilcoxon signed-rank test, p < 0.02). These patterns are not simply due to spatial autocorrelation, as overall Bray-Curtis dissimilarity (from all ASVs) shows no pattern associated with geographic distance across full transects (Figure S6; Kruskall-Wallis rank sum test, p > 0.85).

**Figure 5:**
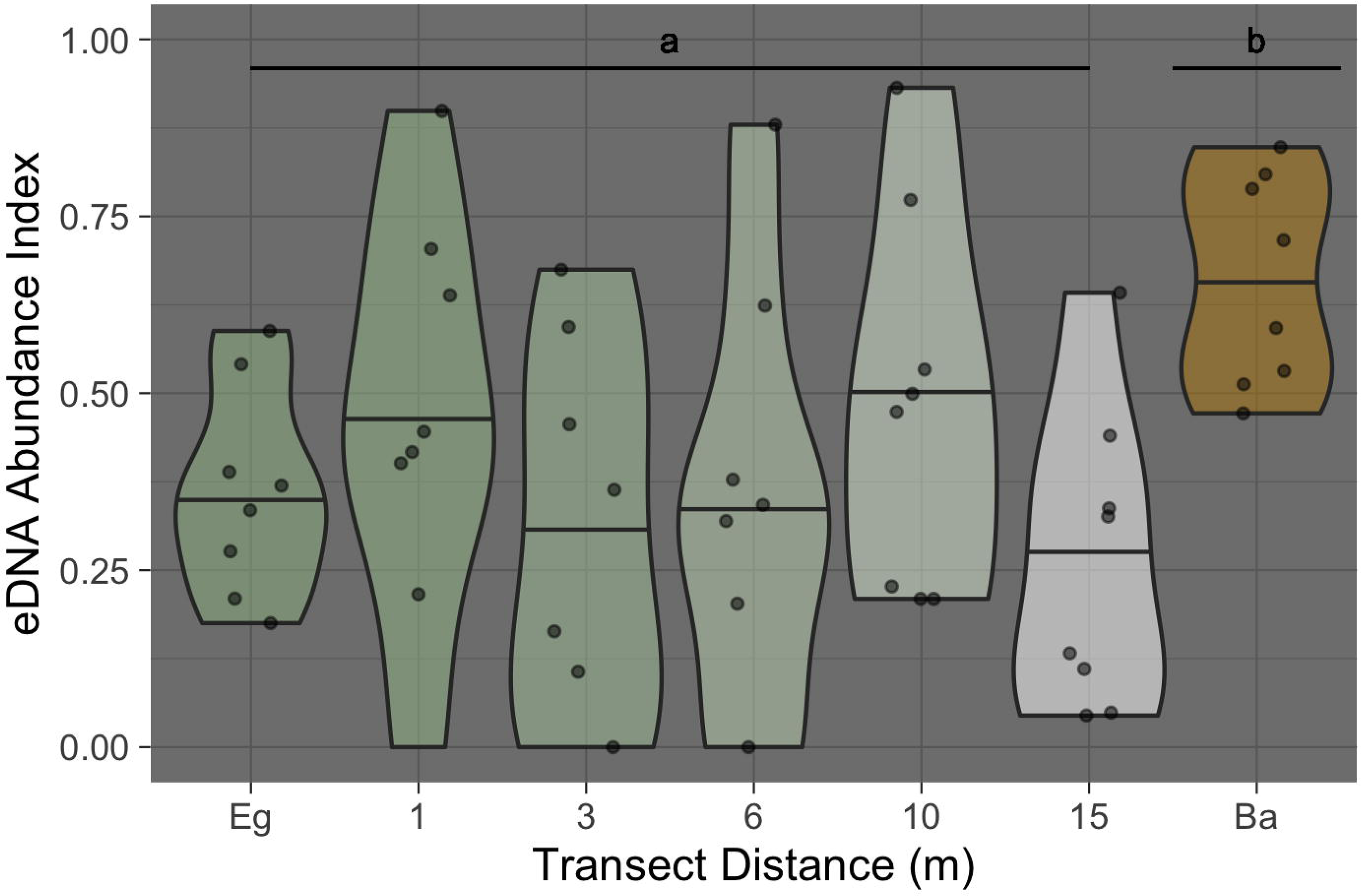
Dinoflagellate eDNA abundance indices plotted for all sites and months combined at each point along the transect from eelgrass (located 47-90 meters inside the edge of the contiguous beds) to bare substrate (located 16-670m outside the edge of the contiguous beds). The central zero-line indicates no bias in abundance between habitat types. The violins are scaled to a fixed maximum width, and display the density of points within the interquartile range (shown by the violin area) around the median (central horizontal line).

## 4 DISCUSSION

In a broad-spectrum eDNA survey of the organisms living in and near to eelgrass, we track the relative abundance of a diverse group of taxa representing thirteen phyla. We demonstrate the ability of eDNA to distinguish communities represented in samples taken only meters apart, and to reveal a significant axis of variance based on proximity to habitat type, despite strong influences of geography and time across sampling events. One major and significant pattern emerges in our analysis: dinoflagellate taxa are more common over bare substrate than within eelgrass beds when highly-abundant, and this putative effect of eelgrass habitat extends at least 15m beyond the edge of the contiguous beds themselves. Because ours was an observational field study, rather than an experiment, we cannot rigorously distinguish among plausible mechanisms for the observed dinoflagellate distributions. Instead, we use the patterns in our own data as well as the relevant scientific literature to evaluate a number of potential hypotheses.

One plausible mechanism is that of an ecological edge effect acting in nearshore zones at the interface between eelgrass and non-eelgrass habitat. “Edge effects” are changes in the distribution or abundance of a species at a boundary between habitats (defined in Boström et al. (5) and see, e.g., Ries and Sisk (43); Macreadie et al. (29)), or in a broader sense, these same effects across many species, observed at a community level. As patch size in a habitat decreases, edge effects become increasingly important factors influencing the distribution of a species. Here, we observe dinoflagellates at increased abundances in sites with bare substrate 16-670 m away from the edge of contiguous eelgrass beds, and not closer (1-15 m). If dinoflagellates were preferentially living at habitat edges, we would expect higher abundance of these taxa at intermediate distances. If, by contrast, the dinoflagellate pattern arose from a community-wide set of interactions in which eelgrass- and non-eelgrass-associated species overlap at habitat edges, we would expect to see community richness peak at or near the habitat boundary. We observe neither of these patterns in our data (Figure 5; Figure S9).

A second plausible mechanism is that slower flow-rates cause dinoflagellate deposition within eelgrass beds, but not outside. Here, too, we would expect to see a continuum of dinoflagellate abundance as a function of eelgrass thinning with distance from the contiguous beds, and we would expect this to be universal among planktonic species of roughly the same size. Instead, we observe a halo of lowered dinoflagellate abundance even when shoot densities are a few per square meter or lower, and do not see this same pattern for other single-celled algae such as taxa within the groups chlorophyta or bacillariophyta. Additionally, we recover eDNA from multiple benthic families (for example, Dendrasteridae, Tellinidae, and Veneridae). It therefore seems unlikely that physical factors driving particle deposition alone produce the observed pattern.

A third possibility is that predatory taxa exist in greater abundance within eelgrass beds and thereby consume dinoflagellates in larger quantities within this habitat. This mechanism appears unlikely for two reasons. First, many organisms that eat dinoflagellates also consume diatoms and other single-celled algae; as stated above, we do not observe the same patterns of reduced abundance in and around eelgrass beds for these other taxa. Additionally, a test of habitat associations at the family level reveals no predators of dinoflagellates with significantly higher abundance in eelgrass habitat relative to bare substrate.

We find greater support for the hypothesis that allelopathy from within the eelgrass bed excludes dinoflagellates. A specific allelopathy against microalgal species by *Z. marina* was first described over 30 years ago (17). More recent evidence suggests this negative interaction applies to multiple HAB taxa that cause paralytic or diarrhetic shellfish poisoning (including *Alexandrium*, a genus observed in this study), and is mediated locally by a variety of strains of eelgrass-associated algicidal and growth-inhibiting bacteria, particularly from *Erythrobacter, Teredinibacter, Gaetbulibacter*, and *Arthrobacter* (21) (though the eDNA primers employed here amplify eukaryotes almost exclusively and therefore do not allow us to test this mechanism). However, in our dataset the repressive effect of eelgrass notably does not extend at the phylum level to other phytoplankton such as diatoms (*Bacillariophyta*) and green algae (*Chlorophyta*), despite reports that *Z. marina* habitat can deter members of these taxa as well (reviewed in Gross (15)). If we are witnessing patterns associated with an allelopathic interaction between microalgae and eelgrass mediated by bacterial species, it is possible that local variation in the *Z. marina* microbiome (such as that described in Bengtsson et al. (4)) could produce disparate patterns for various microalgal community members. Regardless of mechanism, the taxonomically broad pattern of lowered dinoflagellate abundance within contiguous eelgrass beds and in a halo of influence up to 15 m in radius surrounding the habitat demands explanation.

Dinoflagellates with consistent patterns of abundance decrease within and around eelgrass habitat in our dataset include species from the genera *Heterocapsa, Alexandrium, Karlodinium, Protoceratium, Gymnodinium* and unknown taxa from the Kareniaceae family, each of which have at least one member included in local microscopy-based monitoring programs (53; 2); our eDNA methodology thus agrees broadly with previous visual identification of microalgae in this region. Of particular interest are dinoflagellate taxa that include HAB-forming members: the resident species of *Alexandrium* (*A. catanella*) causes paralytic shellfish poisoning via production of saxitoxin (STX; Wiese et al. (55)), species from the genus *Protoceratium* (e.g. *P. reticulatum*) produce yessotoxins (YTXs), whose effects on human consumers of contaminated shellfish are complex and unclear (reviewed in Tubaro et al. (52)), and some species within the family Kareniaceae produce ichthyotoxic karlotoxins (Bachvaroff et al. (3)). Saxotoxin, yessotoxins, and karlotoxins impact aquaculture and harvest industries directly; detection of STX at concentrations greater than 80 *μ*g STXequiv/100 g is routinely responsible for regional harvest closures (31), shellfish containing more than 0.1 *μ*g YTX equiv/100g may not be sold to markets within the European Union, although this toxin is not currently regulated within the US (50), and karlotoxins can cause millions of dollars in losses for fisheries in single bloom events (16). In summary, the dinoflagellate taxa deterred by eelgrass habitat in this study have high relevance for local shellfish management decisions, particularly as HABs (including *Alexandrium*) are intensifying with recent ocean warming in the North Pacific (13), and are associated with an increase in the number of shellfish harvesting closures (51; 31).

In order to understand the relationship of *Z. marina* to ecosystem and human health, as well as to shellfish farming and harvest, it is critical to consider our addition of a possible ‘action-at-a-distance’ element to the existing eelgrass-dinoflagellate interaction model. Given the protected status of *Z. marina* habitat on the Pacific Coast of the United States, the goals of the shellfish industry and eelgrass conservation are often perceived as being in conflict (9) and policies prohibit shellfish farming and harvesting within or near beds. For example, in Washington State, required buffer zones between shellfish aquaculture and eelgrass range from 3 to 8 m, depending on the agency involved (32). However, our work demonstrates that *Z. marina* habitat may have a protective effect against harmful dinoflagellates within these buffer zones, reducing the potential for shellfish to accumulate HAB toxins from the surrounding waters. Likewise, filter feeders can mitigate microbial disease in adjacent environments, and *Magallana* (née *Crassostrea*) *gigas*, in particular, has recently been shown to lessen the effects of Eelgrass Wasting Disease (EWD) on *Z. marina* (14). As others have begun to suggest, then, eelgrass and oysters may provide critical support to one another in changing marine ecosystems worldwide. Future work will examine their multi-faceted mutualism, particularly in characterizing the taxonomic breadth of potential seagrass and shellfish partnerships, as well as in defining the molecular mechanisms underlying the roles of both beneficial and detrimental microbial intermediates.

## Notes

https://github.com/invertdna/EelgrassHalo

