## Supplemental Material for "A Halo of Reduced Dinoflagellate Abundances In and Around Eelgrass Beds"

### Supplementary tables and figures

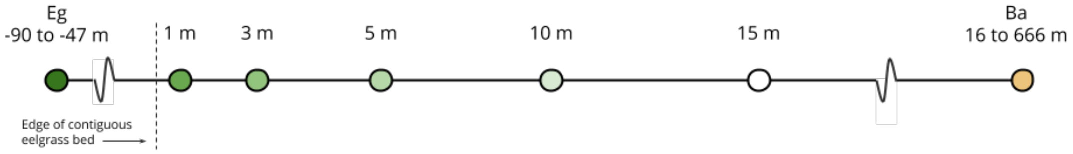


*Figure S1: Schematic of transects from contiguous eelgrass beds (Eg, dark green) to bare substrate (Ba, brown). Distance from the edge of the contiguous eelgrass beds are given in meters; sites outside of the contiguous beds have an average shoot density of less than three shoots per square meter and the bare substrate site is devoid of eelgrass.*

*Table S1: Sample information. For each site, Case Inlet (CI), Port Gamble (PG), Nisqually Reach (NR), Skokomish (SK), and Willapa Bay (WB), approximate transect positions are recorded, as well as latitude, longitude, and the approximate geographic distance of each sample from the eelgrass bed edge, calculated from coordinates. Negative distances indicate samples within the eelgrass bed itself. Columns named May, July, and August list the number of technical replicates passing quality control measures of three sequenced from each bottle of water. NA indicates samples that were not gathered, and asterisks indicate samples for which three technical replicates were sequenced on two separate MiSeq runs to characterize the importance of sequencing run in explaining variation among samples.*

| Site | Position | long | lat | Distance | May | July | August |
|---|---|---|---|---|---|---|---|
| CI | Eelgrass | -122.79645 | 47.358439 | -47 | 3 | 6* | 3 |
| CI | Along 1 | -122.79584 | 47.358455 | 1 | 3 | 3 | 3 |
| CI | Along 3 | -122.796038 | 47.358565 | 3 | 3 | 3 | 3 |
| CI | Along 6 | -122.795971 | 47.358551 | 6 | 3 | 3 | 2 |
| CI | Along 10 | -122.795894 | 47.358481 | 10 | 3 | 3 | 2 |
| CI | Along 15 | -122.795817 | 47.358436 | 15 | 1 | 3 | 2 |
| CI | Bare | -122.79576 | 47.357937 | 57 | 3 | 3 | 3 |
| NR | Eelgrass | -122.726752 | 47.101926 | NA | 3 | 3 | 2 |
| NR | Bare | -122.726386 | 47.101713 | NA | 3 | 3 | 3 |
| PG | Eelgrass | -122.58292 | 47.847983 | -80 | 3 | 3 | 3 |
| PG | Along 1 | -122.583221 | 47.84866 | 1 | 3 | 3 | 2 |
| PG | Along 3 | -122.583157 | 47.848705 | 3 | 3 | 2 | 2 |
| PG | Along 6 | -122.583222 | 47.848725 | 6 | 3 | 3 | 0 |
| PG | Along 10 | -122.583278 | 47.848756 | 10 | 3 | 3 | 3 |
| PG | Along 15 | -122.583258 | 47.848781 | 15 | 3 | 3 | 3 |
| PG | Bare | -122.58383 | 47.842676 | 666 | 3 | 6* | 3 |
| SK | Eelgrass | -123.156623 | 47.354332 | -52 | 3 | 3 | 3 |

| | | | | | | | |
|---|---|---|---|---|---|---|---|
| SK | Along 1 | -123.157147 | 47.354626 | 1 | NA | 3 | 3 |
| SK | Along 3 | -123.157132 | 47.354585 | 3 | NA | 3 | 2 |
| SK | Along 6 | -123.157116 | 47.354634 | 6 | NA | 3 | 3 |
| SK | Along 10 | -123.157162 | 47.354644 | 10 | NA | 3 | 3 |
| SK | Along 15 | -123.157185 | 47.354733 | 15 | NA | 3 | 2 |
| SK | Bare | -123.157314 | 47.35502 | 45 | 3 | 3 | 3 |
| WB | Eelgrass | -124.02619 | 46.495137 | -90 | 3 | 3 | NA |
| WB | Along 1 | -124.02622 | 46.494334 | 1 | 3 | 3 | 2 |
| WB | Along 3 | -124.02627 | 46.494347 | 3 | 3 | 3 | 3 |
| WB | Along 6 | -124.02626 | 46.494425 | 6 | 3 | 3 | 3 |
| WB | Along 10 | -124.02624 | 46.494437 | 10 | 3 | 3 | 3 |
| WB | Along 15 | -124.02619 | 46.494479 | 15 | 3 | 3 | 3 |
| WB | Bare | -124.026136 | 46.494479 | 16 | 3 | 3 | 3 |

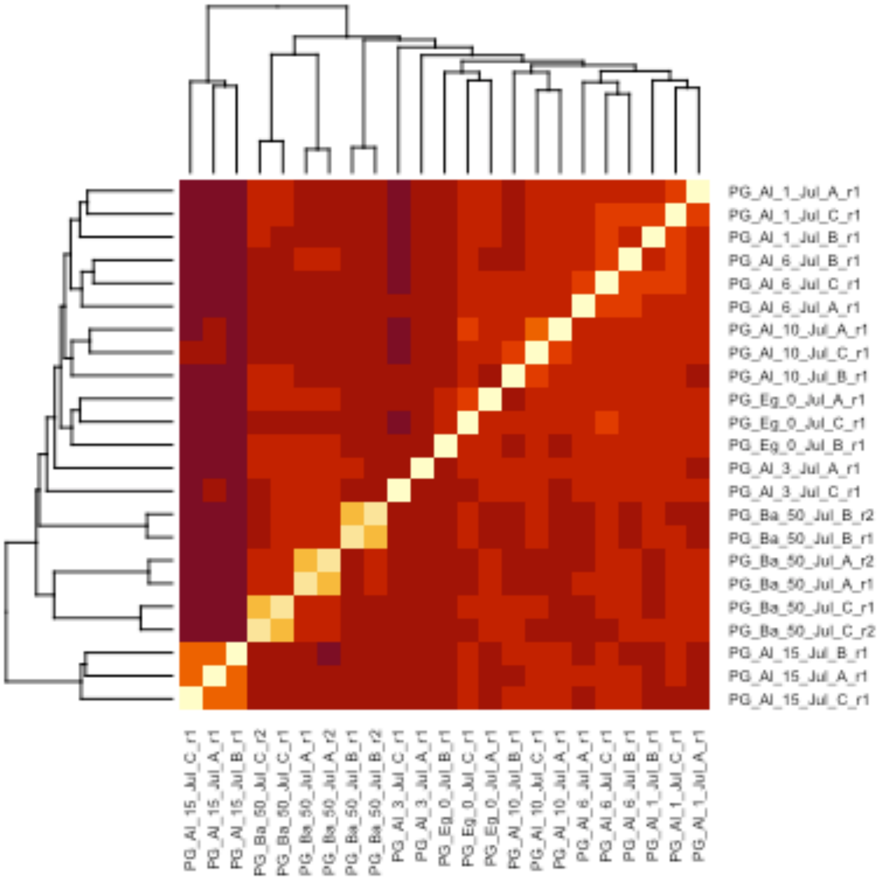

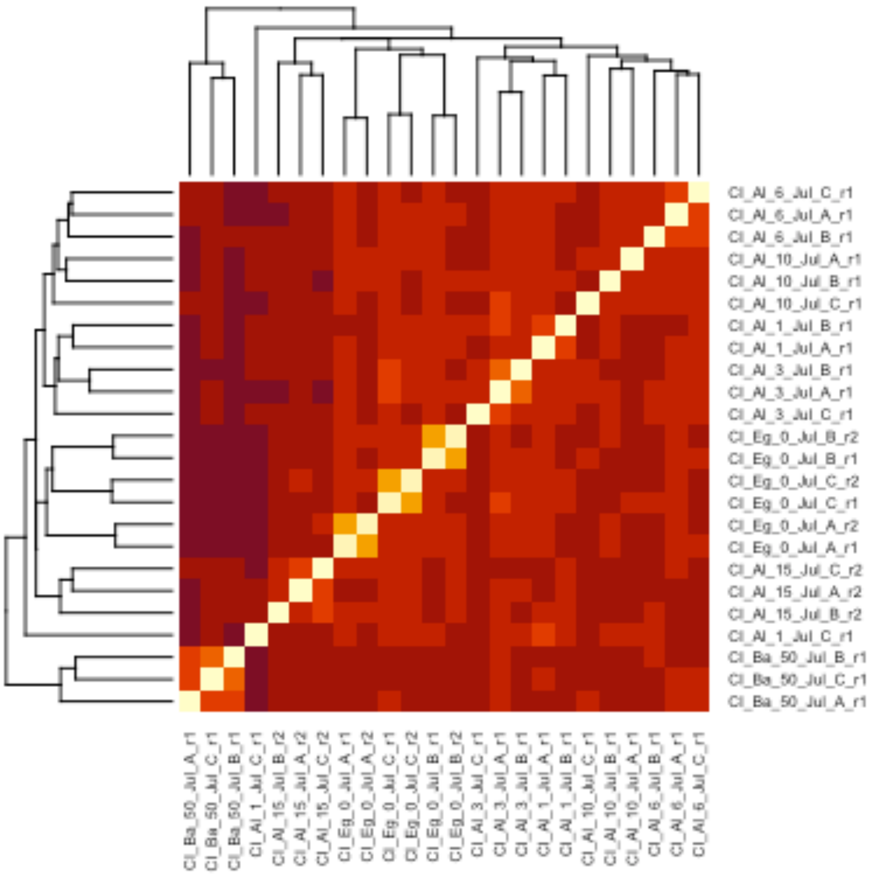


*Figure S2: Hierarchical clustering from transect Bray-Curtis distance matrices in which three technical replicates were sequenced on two different runs. Names of technical replicates contain sample information separated by '_' as follows: Site abbreviation, position abbreviation, transect distance, month, replicate, and sequencing run. Note that all replicates from Miseq run 2 (r2) cluster with the corresponding replicates from Miseq run 1 (r1) for both Port Gamble July and Case Inlet July transects.*

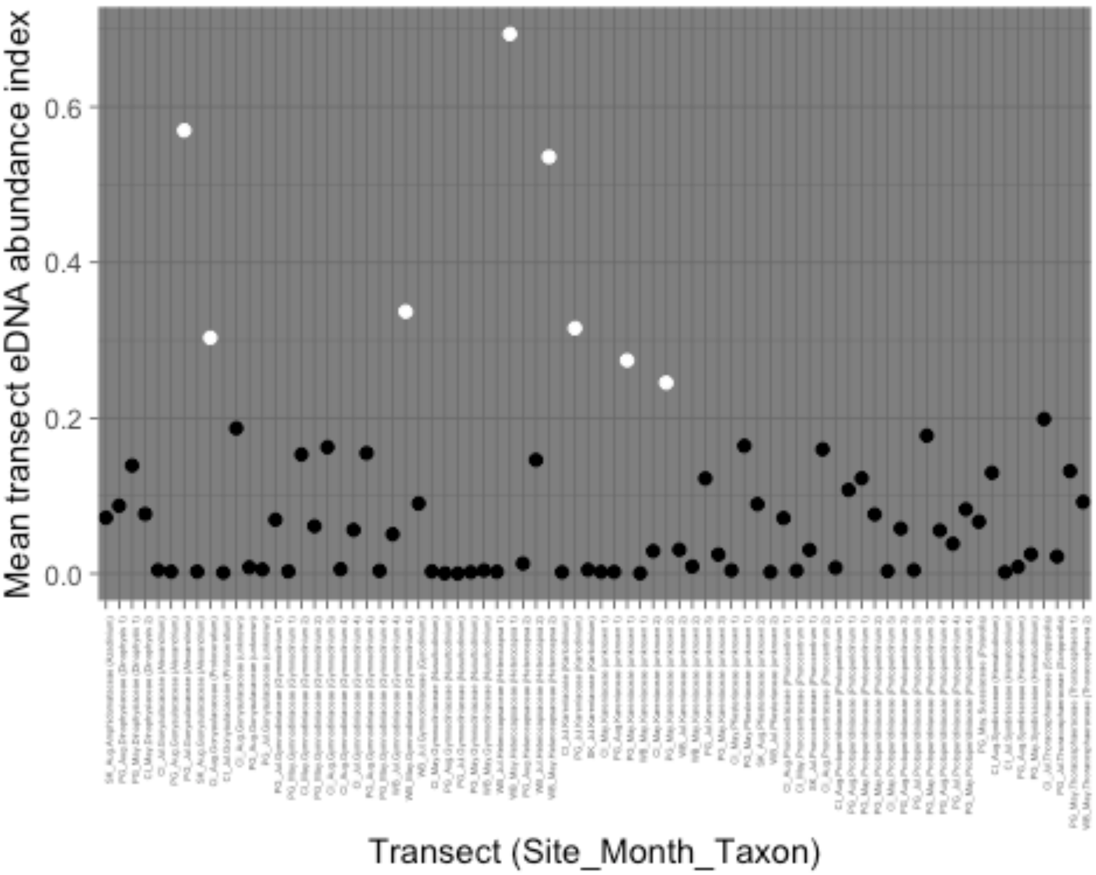


*Figure S5: High- and low-abundance transects generated with kmeans unsupervised machine learning. Black points indicate high-abundance transects; white points indicate low-abundance transects*

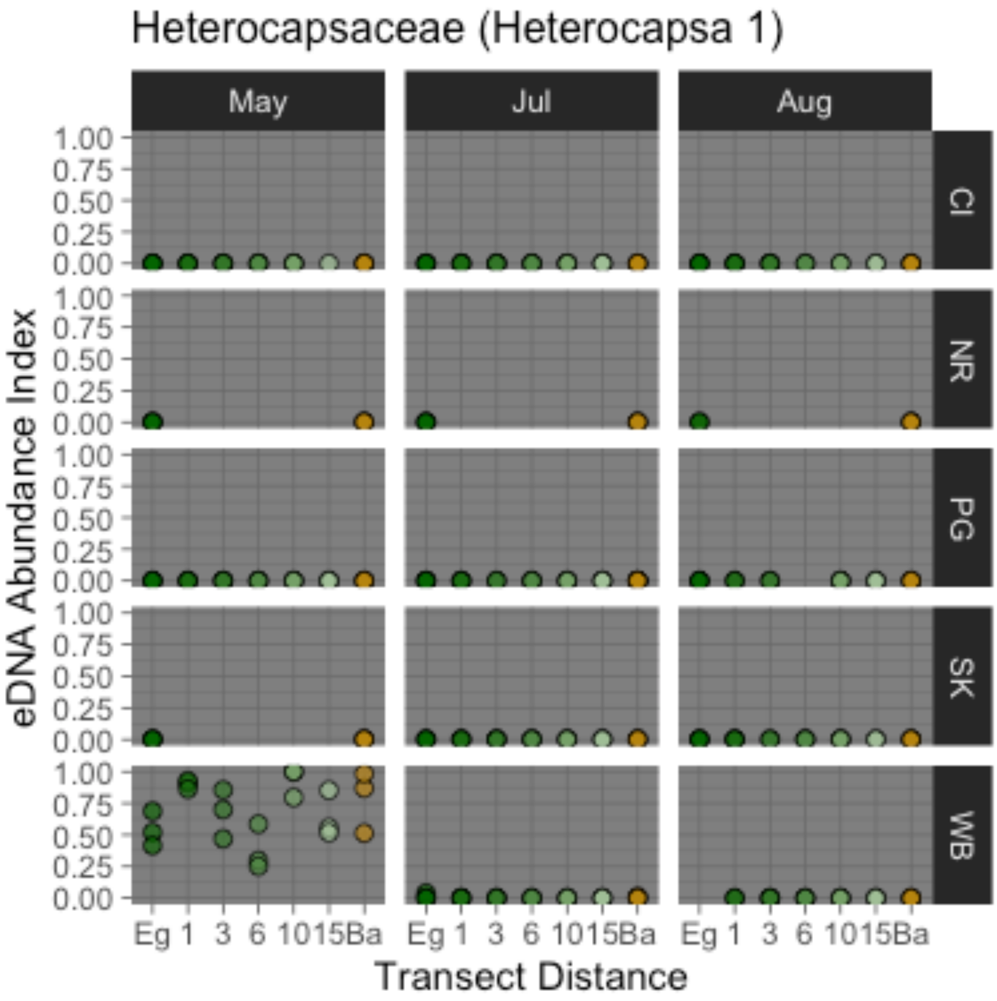
Heterocapsaceae (Heterocapsa 1)
May
Jul
Aug
CI
NR
PG
SK
WB
1.00
0.75
0.50
0.25
0.00
eDNA Abundance Index
Eg 1 3 6 10 15 Ba
Transect Distance

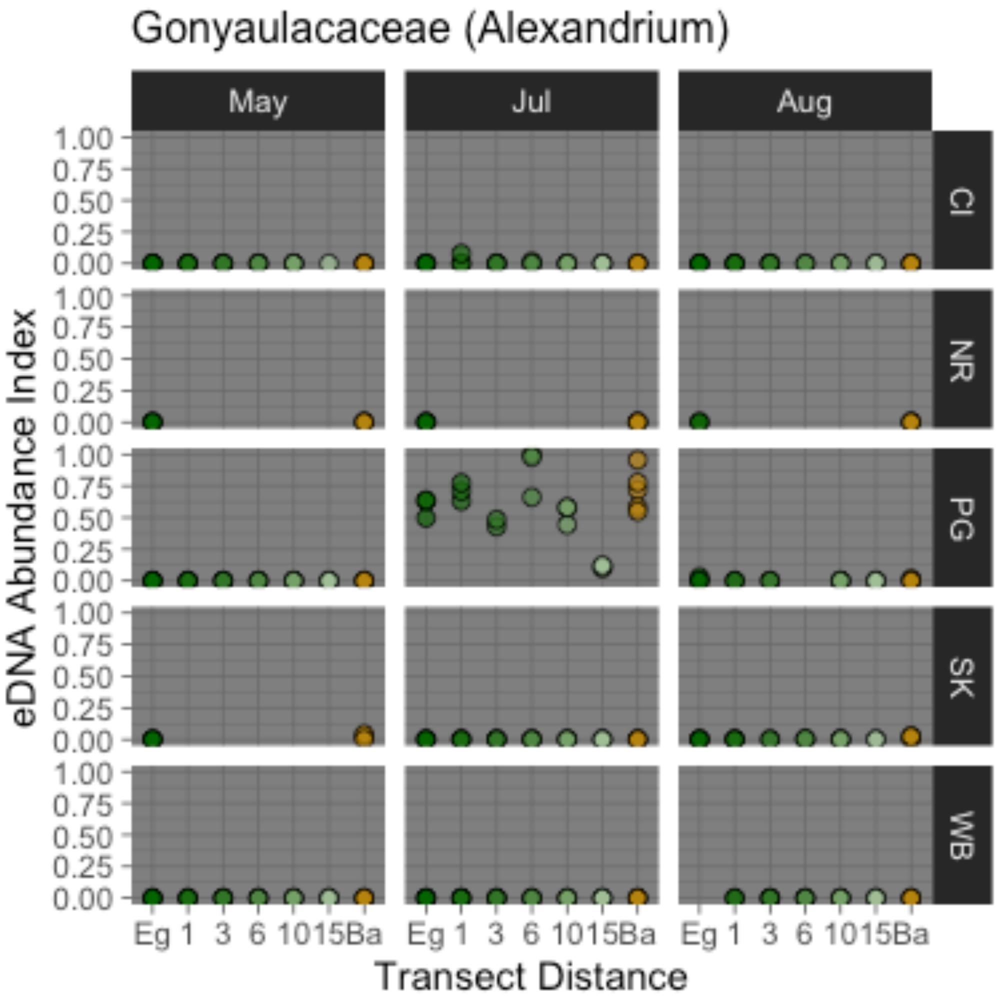
Gonyaulacaceae (Alexandrium)
May
Jul
Aug
CI
NR
PG
SK
WB
1.00
0.75
0.50
0.25
0.00
eDNA Abundance Index
Eg 1 3 6 10 15 Ba
Transect Distance

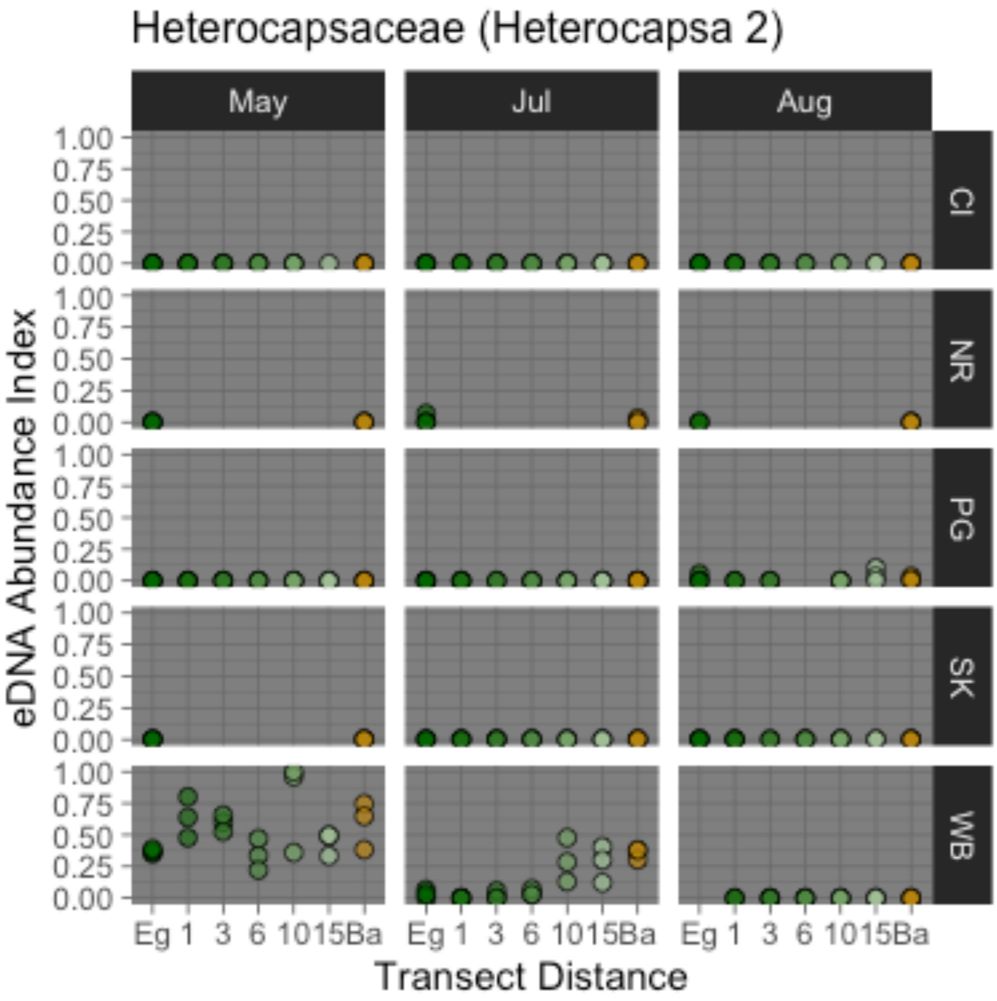
Heterocapsaceae (Heterocapsa 2)
May
Jul
Aug
CI
NR
PG
SK
WB
1.00
0.75
0.50
0.25
0.00
eDNA Abundance Index
Eg 1 3 6 10 15 Ba
Transect Distance

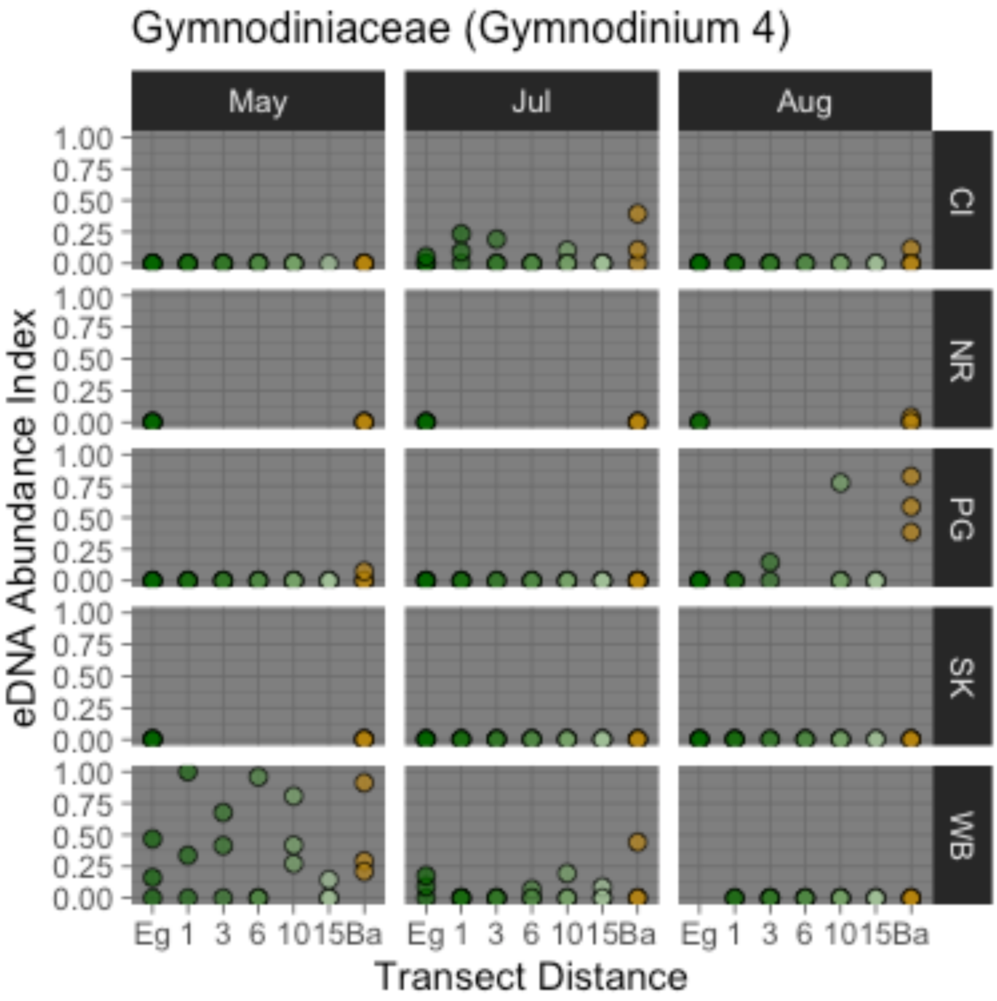

Gymnodiniaceae (Gymnodinium 4)
May
Jul
Aug
CI
NR
PG
SK
WB
1.00
0.75
0.50
0.25
0.00
eDNA Abundance Index
Eg 1 3 6 10 15 Ba
Transect Distance

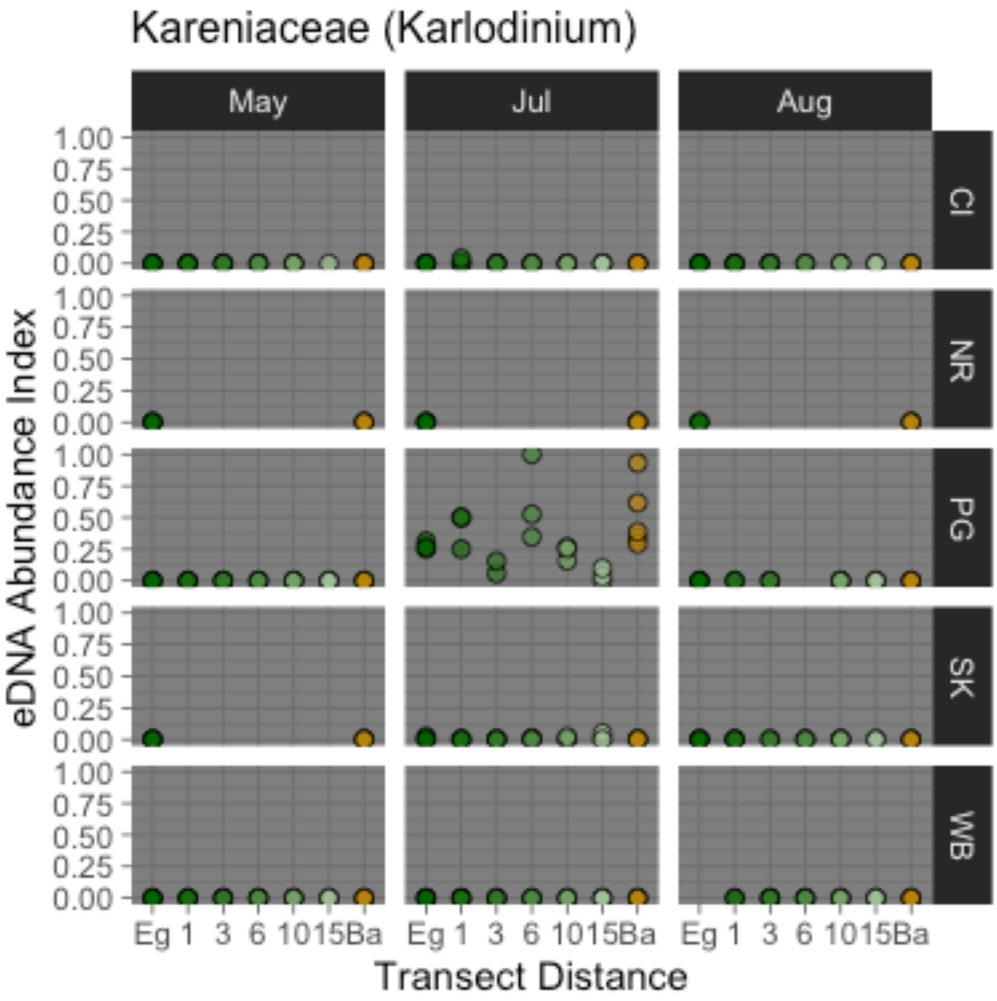

Kareniaceae (Karlodinium)
May
Jul
Aug
CI
NR
PG
SK
WB
1.00
0.75
0.50
0.25
0.00
eDNA Abundance Index
Eg 1 3 6 10 15 Ba
Transect Distance

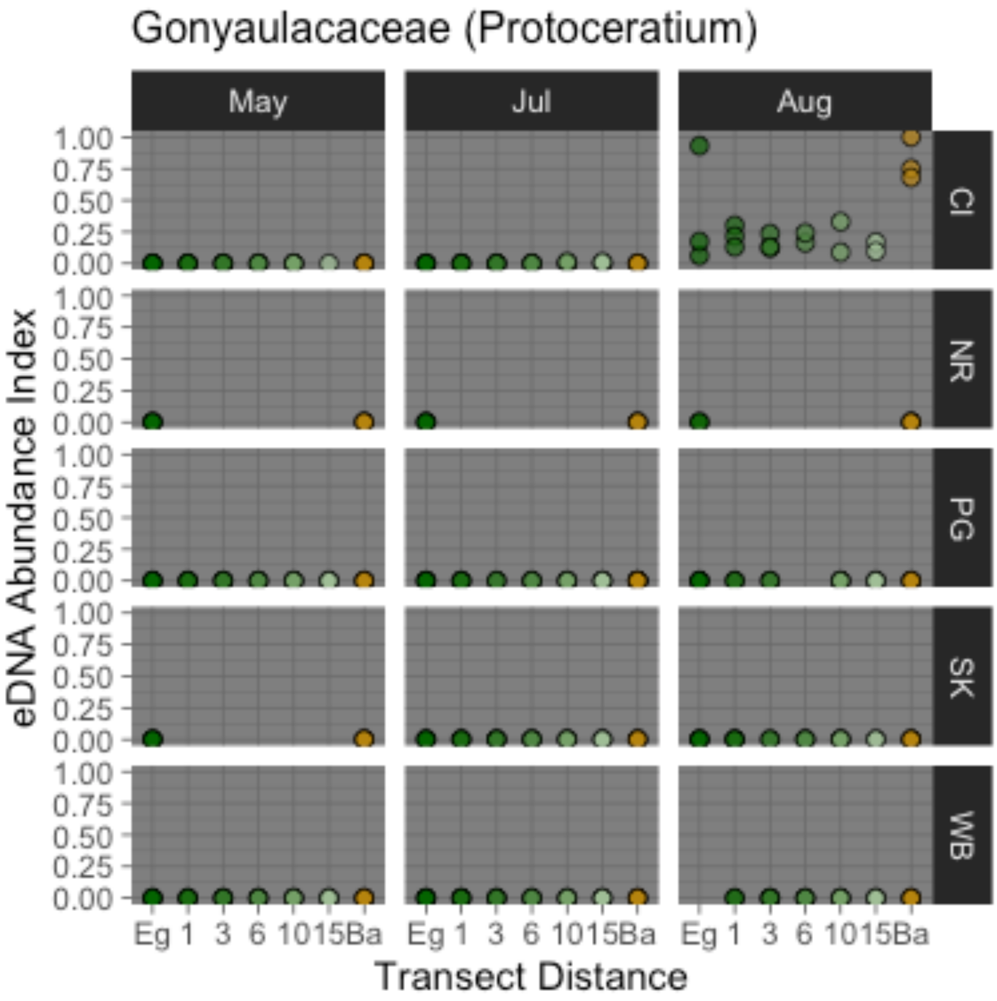

Gonyaulacaceae (Protoceratium)
May
Jul
Aug
CI
NR
PG
SK
WB
1.00
0.75
0.50
0.25
0.00
eDNA Abundance Index
Eg 1 3 6 10 15 Ba
Transect Distance

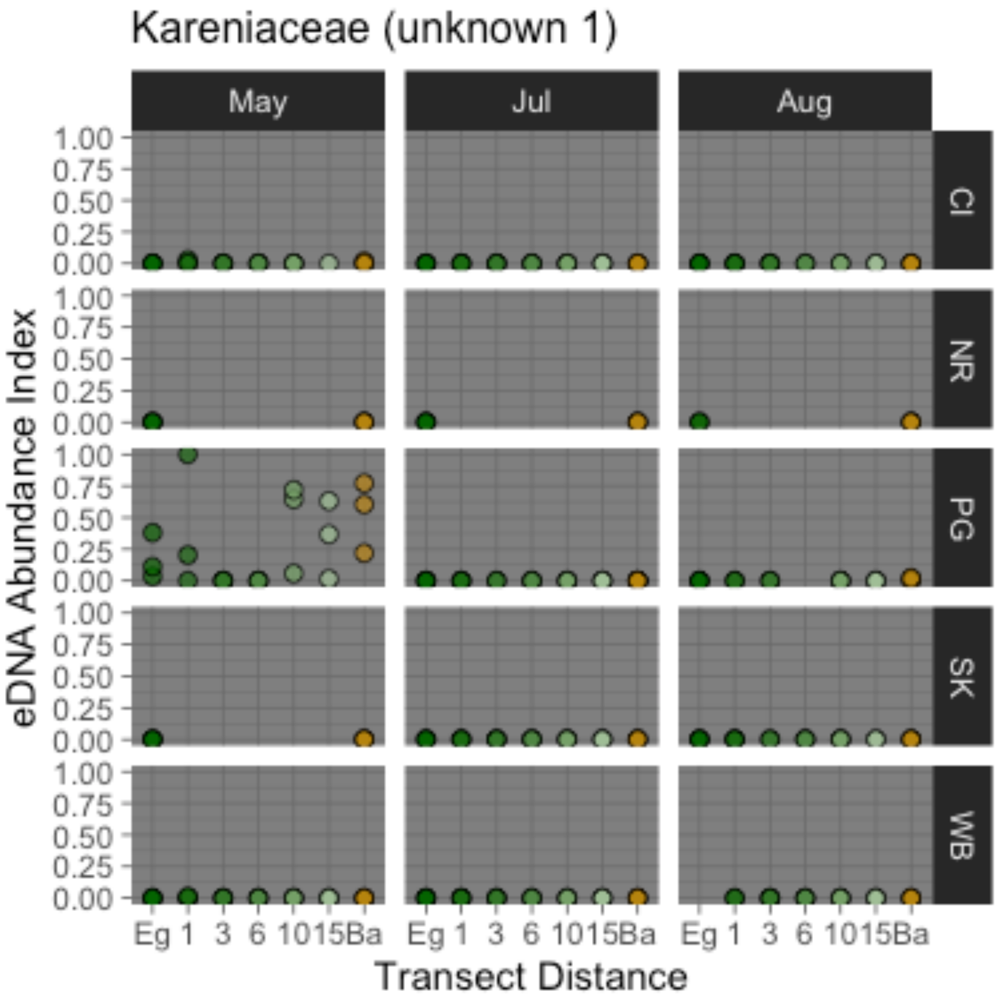

Kareniaceae (unknown 1)
May
Jul
Aug
CI
NR
PG
SK
WB
1.00
0.75
0.50
0.25
0.00
eDNA Abundance Index
Eg 1 3 6 10 15 Ba
Transect Distance

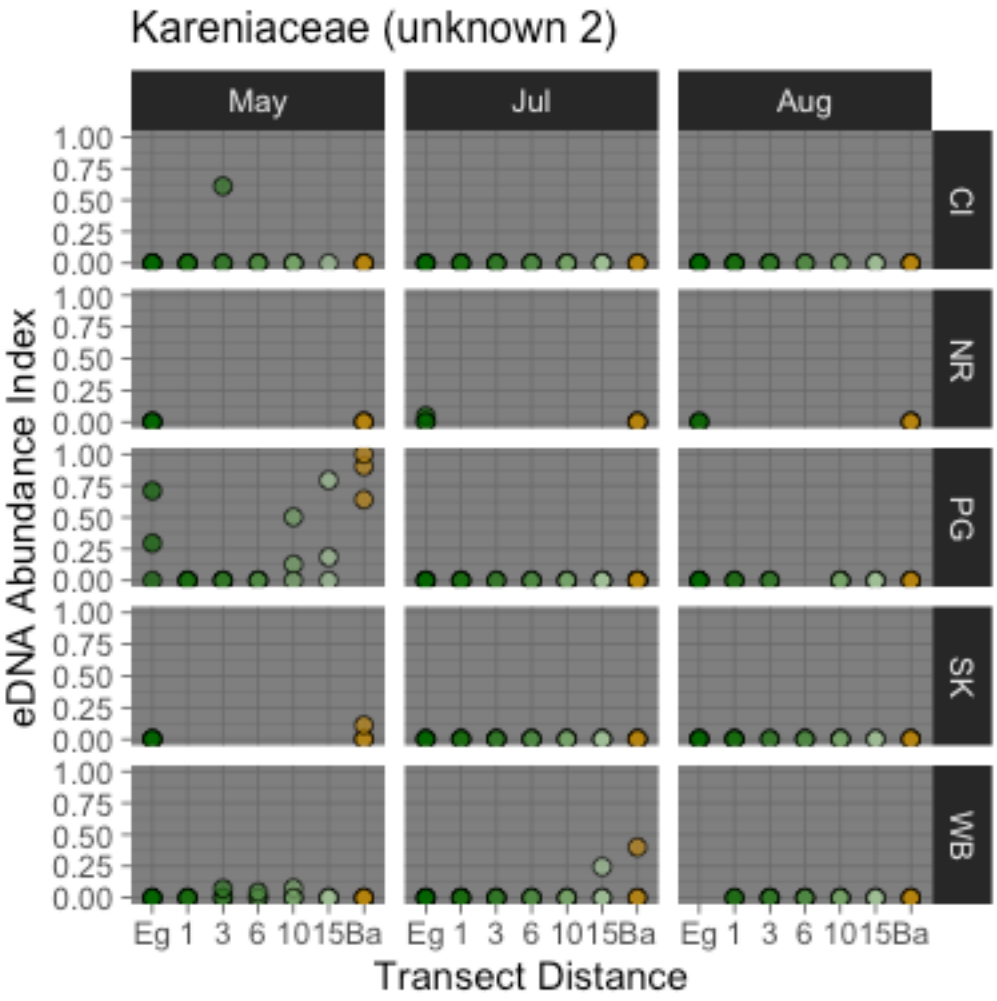


*Figure S3: eDNA indices from each technical replicate of biological samples for each dinoflagellate family represented in our dataset by >1000 sequence reads, plotted at all sites for which complete transect data were available. Color indicates proximity to eelgrass habitat (dark green) versus bare substrate (brown).*

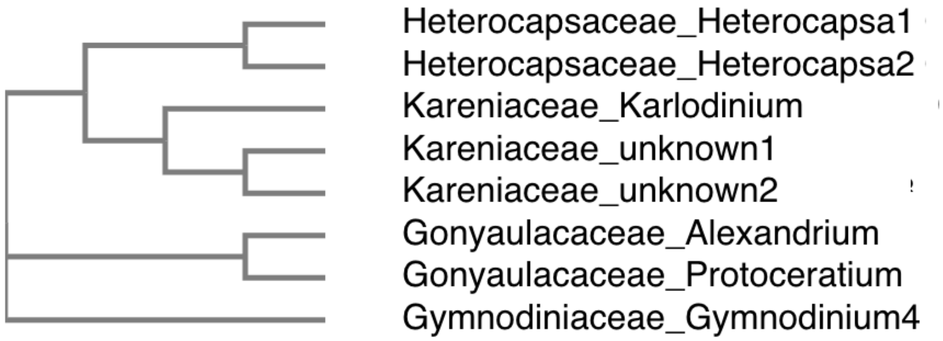


*Figure S5: Phylogeny of dinoflagellate sequences from high-abundance transects. Individual sequences are named with Family_Genus information (when known).*

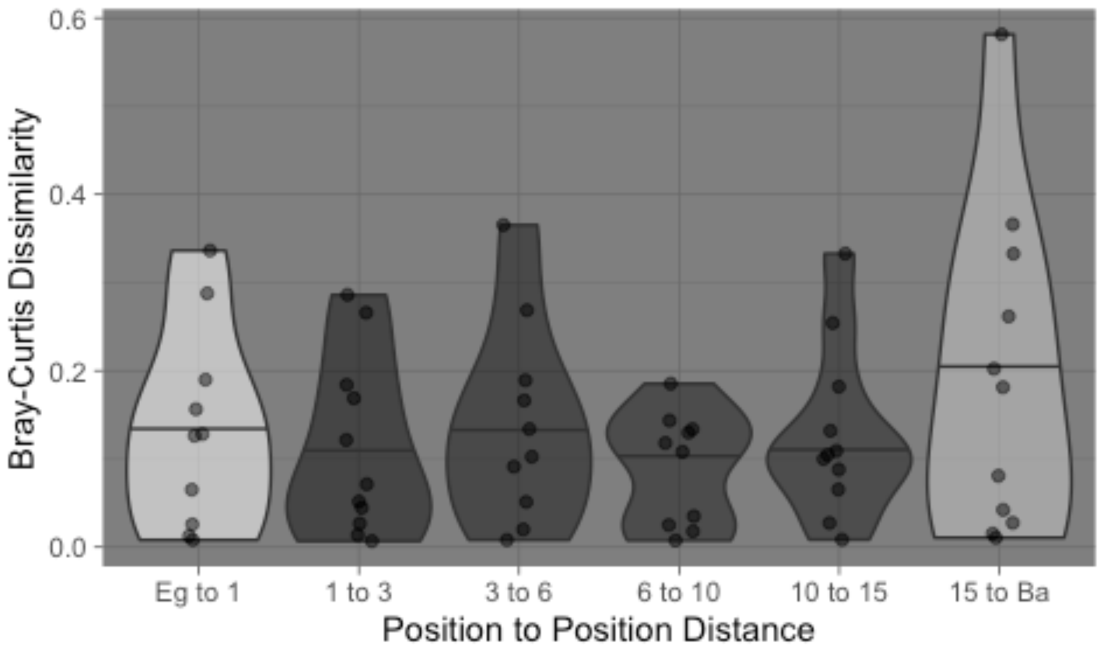


*Figure S6: Bray-Curtis dissimilarity between eDNA communities surveyed at adjacent points along each full transect (all sites and months). Shading of violins indicates median spatial distance between communities (dark = closer, light = more distant).*

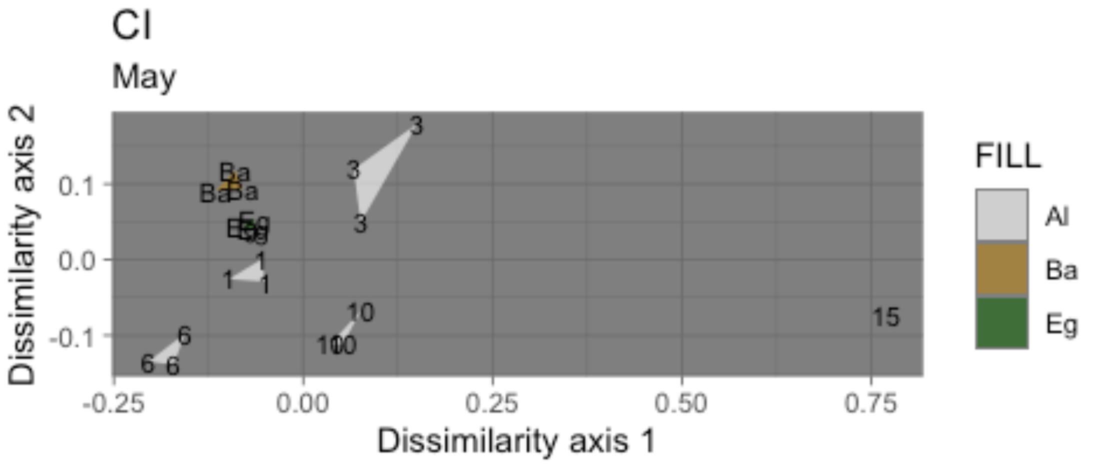

CI
May
Dissimilarity axis 2
Dissimilarity axis 1
0.1
0.0
-0.1
-0.25
0.00
0.25
0.50
0.75
FILL
Al
Ba
Eg

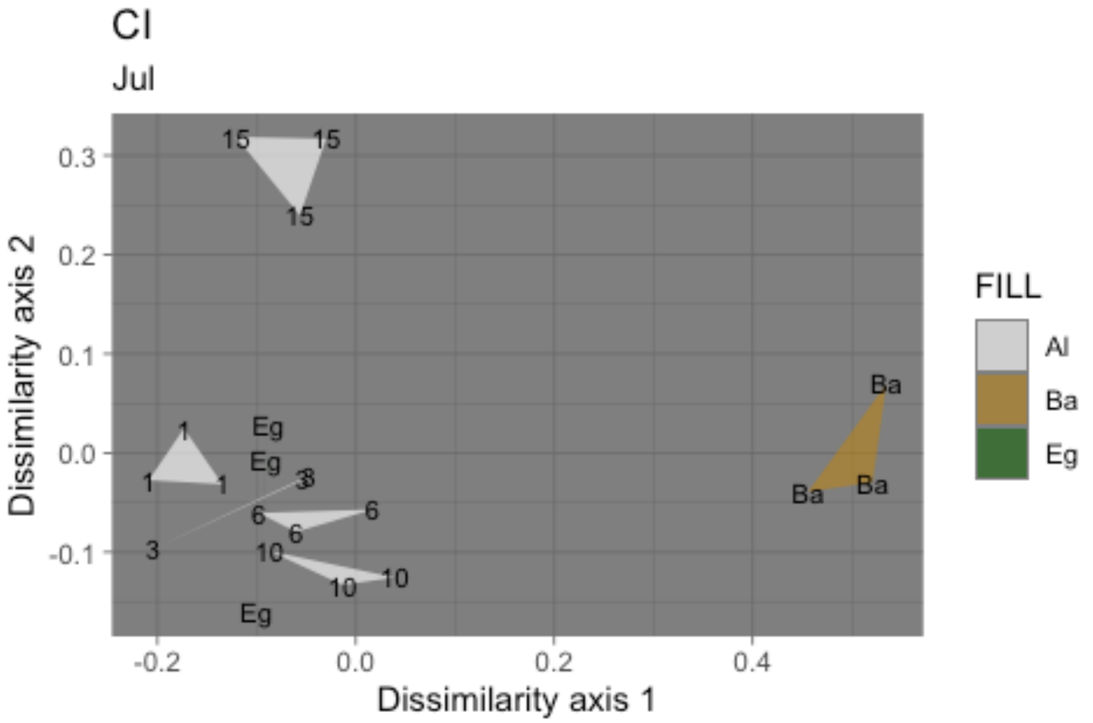

Cl
Jul
Dissimilarity axis 2
Dissimilarity axis 1
0.3
0.2
0.1
0.0
-0.1
-0.2
0.0
0.2
0.4
15
15
15
1
1
1
Eg
Eg
Eg
3
3
3
6
6
10
10
10
Ba
Ba
Ba
FILL
Al
Ba
Eg

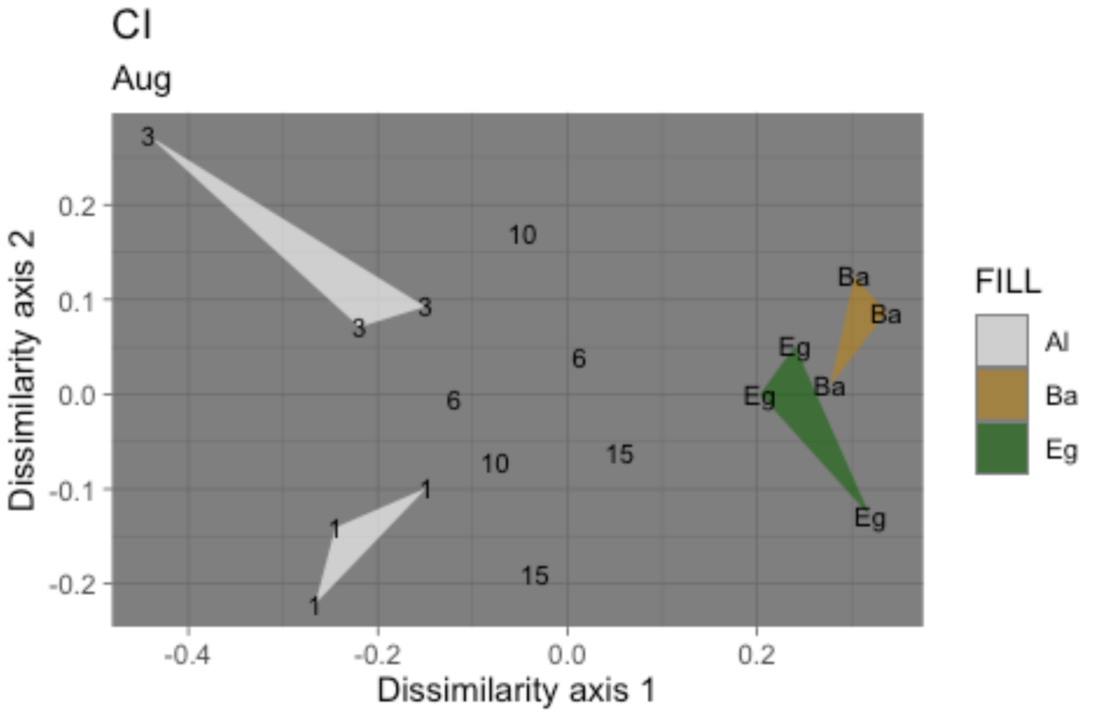

Cl
Aug
Dissimilarity axis 2
Dissimilarity axis 1
0.2
0.1
0.0
-0.1
-0.2
-0.4
-0.2
0.0
0.2
3
3
3
10
6
6
10
15
15
1
1
1
Ba
Ba
Ba
Eg
Eg
Eg
FILL
Al
Ba
Eg

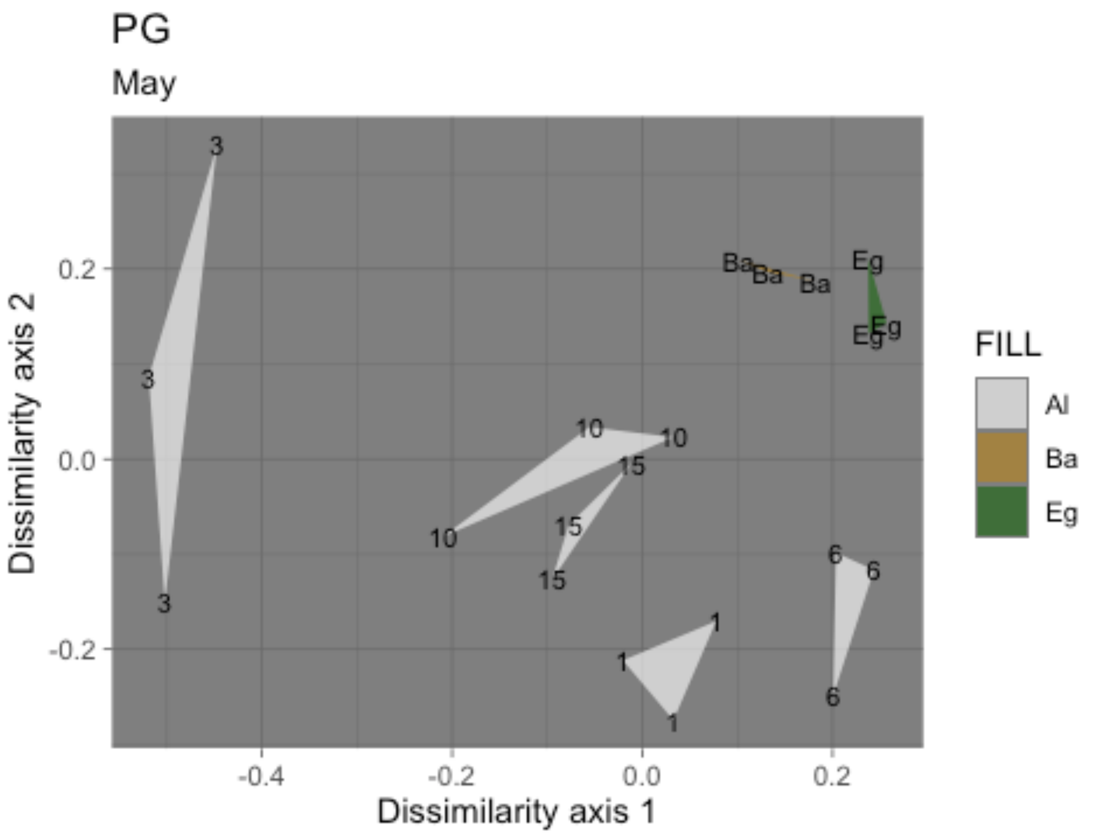
PG
May
Dissimilarity axis 2
Dissimilarity axis 1
0.2
0.0
-0.2
-0.4
-0.2
0.0
0.2
FILL
Al
Ba
Eg

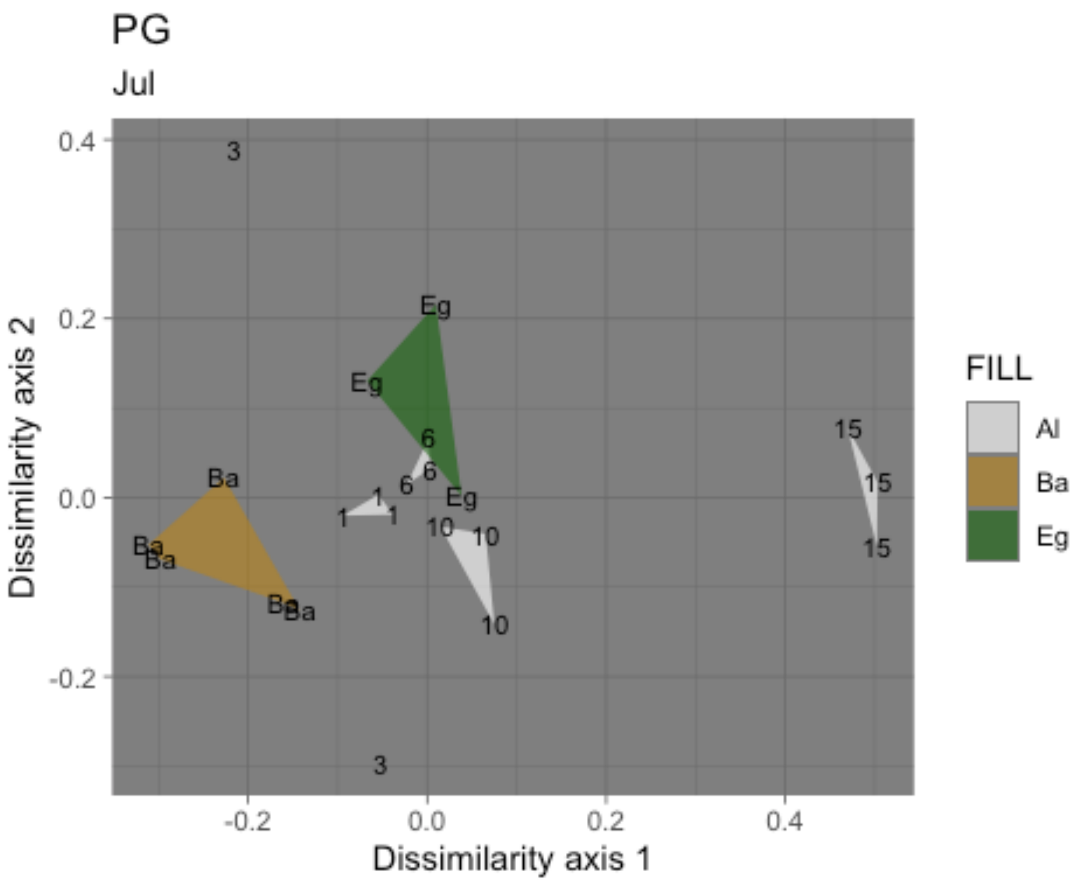
PG
Jul
Dissimilarity axis 2
Dissimilarity axis 1
0.4
0.2
0.0
-0.2
-0.2
0.0
0.2
0.4
3
Eg
Eg
6
6
6
Eg
1
1
1
10
10
10
Ba
Ba
Ba
Ba
Ba
15
15
15
3
FILL
Al
Ba
Eg

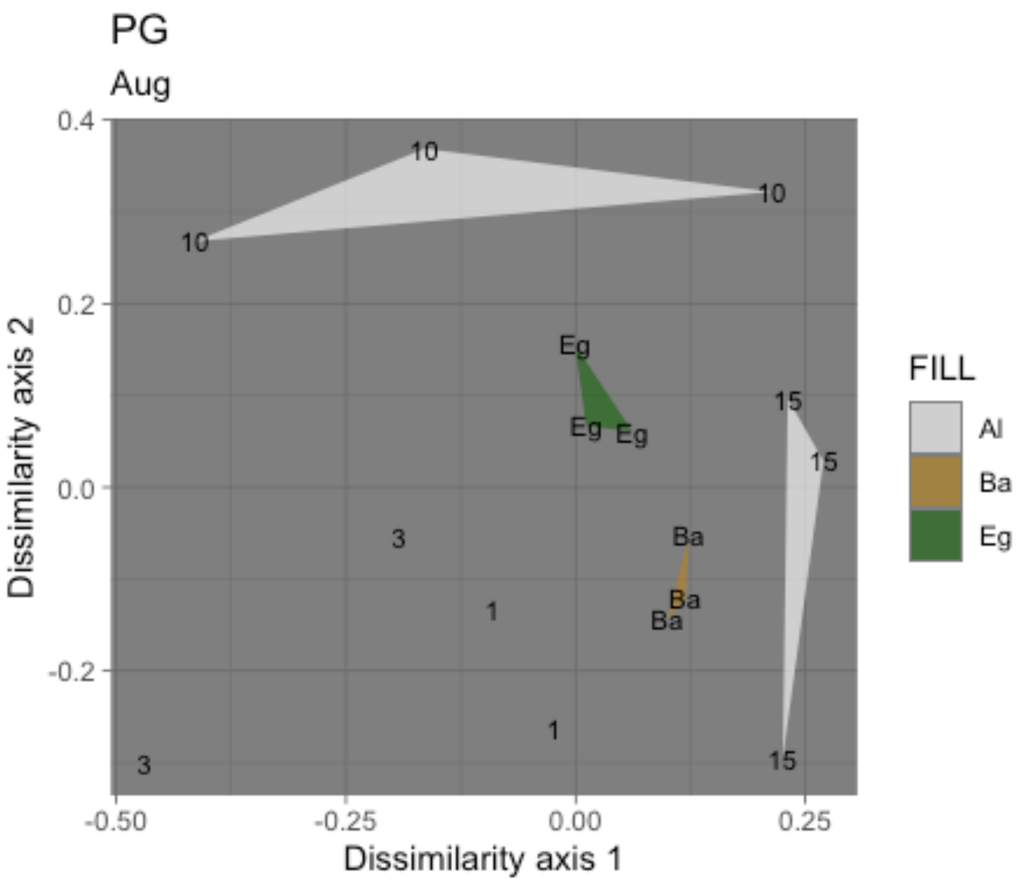

PG
Aug
0.4
0.2
0.0
-0.2
-0.50
-0.25
0.00
0.25
Dissimilarity axis 1
Dissimilarity axis 2
10
10
10
Eg
Eg
Eg
15
15
15
Ba
Ba
Ba
3
1
1
3
FILL
Al
Ba
Eg

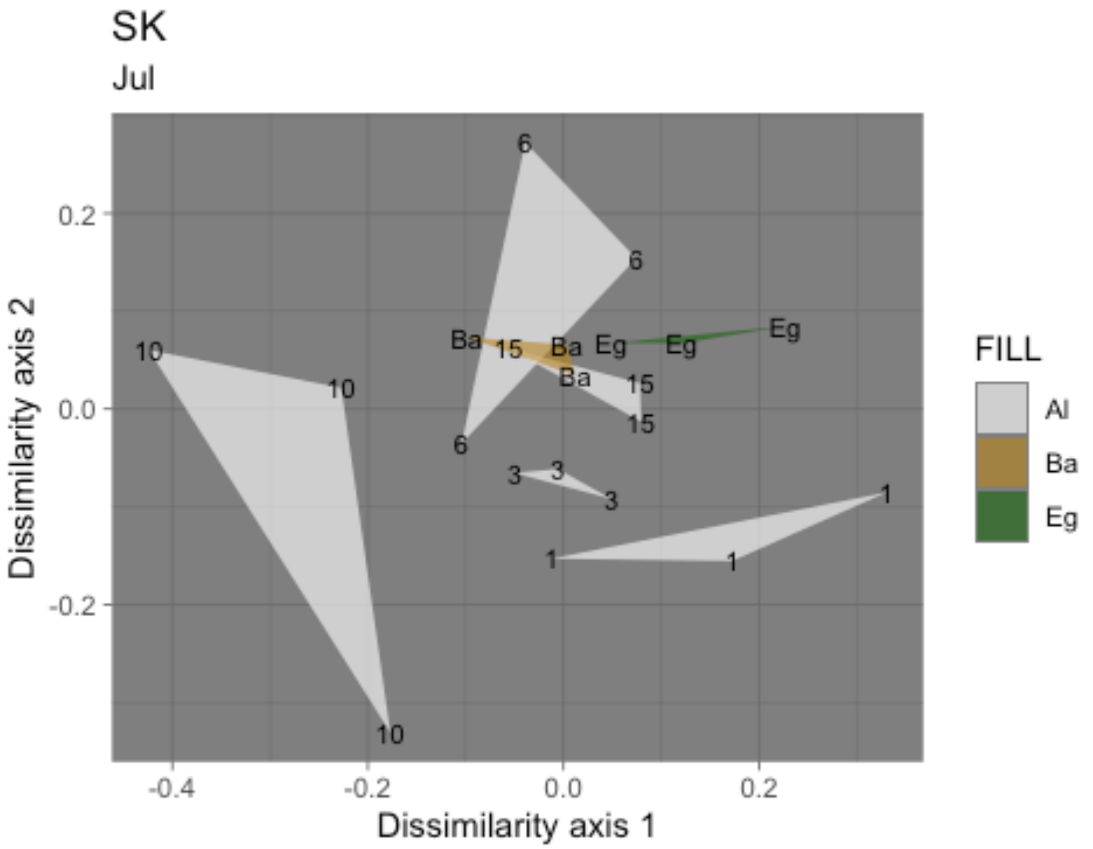
SK
Jul
Dissimilarity axis 2
Dissimilarity axis 1
0.2
0.0
-0.2
-0.4
-0.2
0.0
0.2
6
6
Ba
15
Ba
Eg
Eg
Eg
Ba
15
15
10
10
10
6
3
3
3
1
1
1
FILL
Al
Ba
Eg

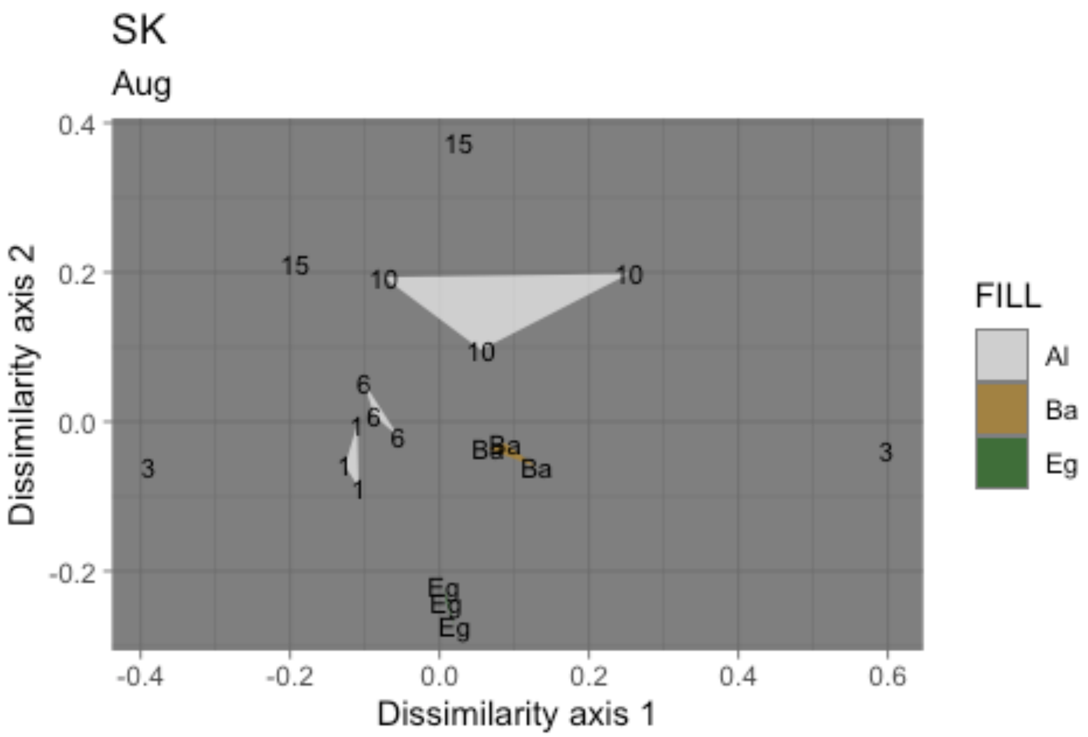

SK
Aug
Dissimilarity axis 2
Dissimilarity axis 1
0.4
0.2
0.0
-0.2
-0.4
-0.2
0.0
0.2
0.4
0.6
15
15
10
10
10
6
6
6
1
1
1
3
3
Ba
Ba
Ba
Eg
Eg
Eg
FILL
Al
Ba
Eg

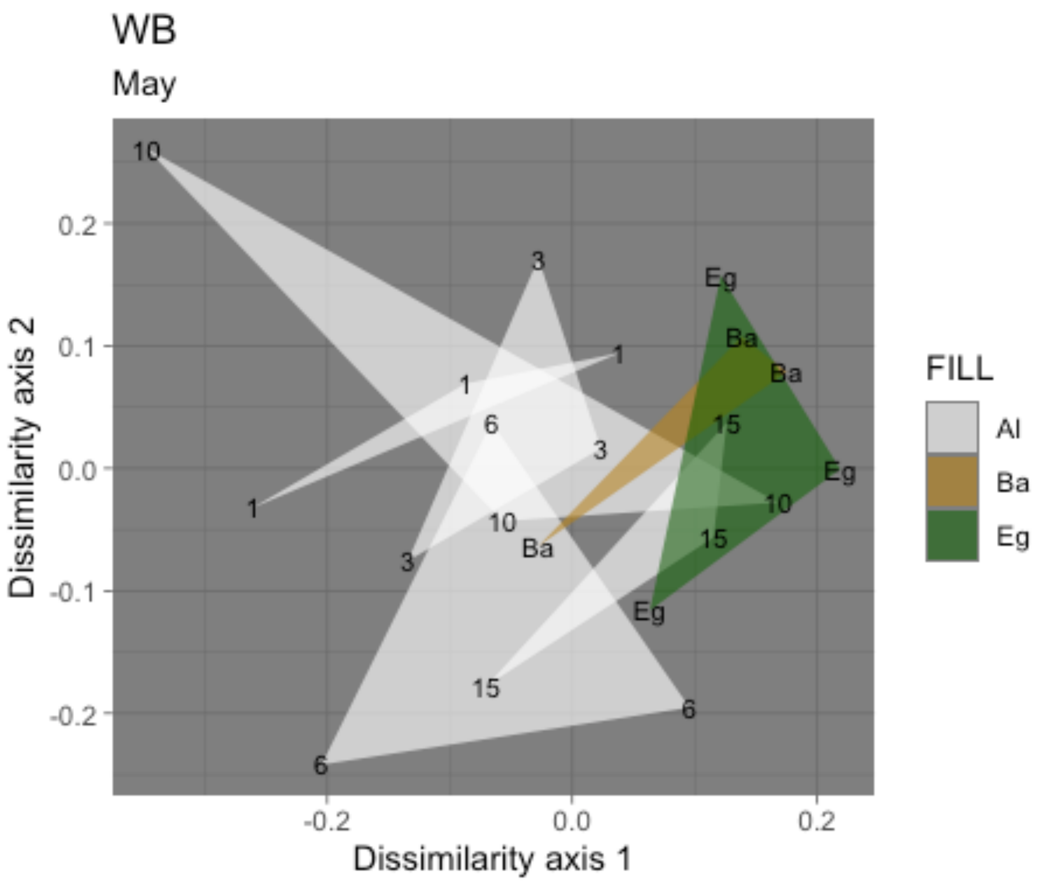

WB
May
Dissimilarity axis 2
Dissimilarity axis 1
0.2
0.1
0.0
-0.1
-0.2
-0.2
0.0
0.2
FILL
Al
Ba
Eg

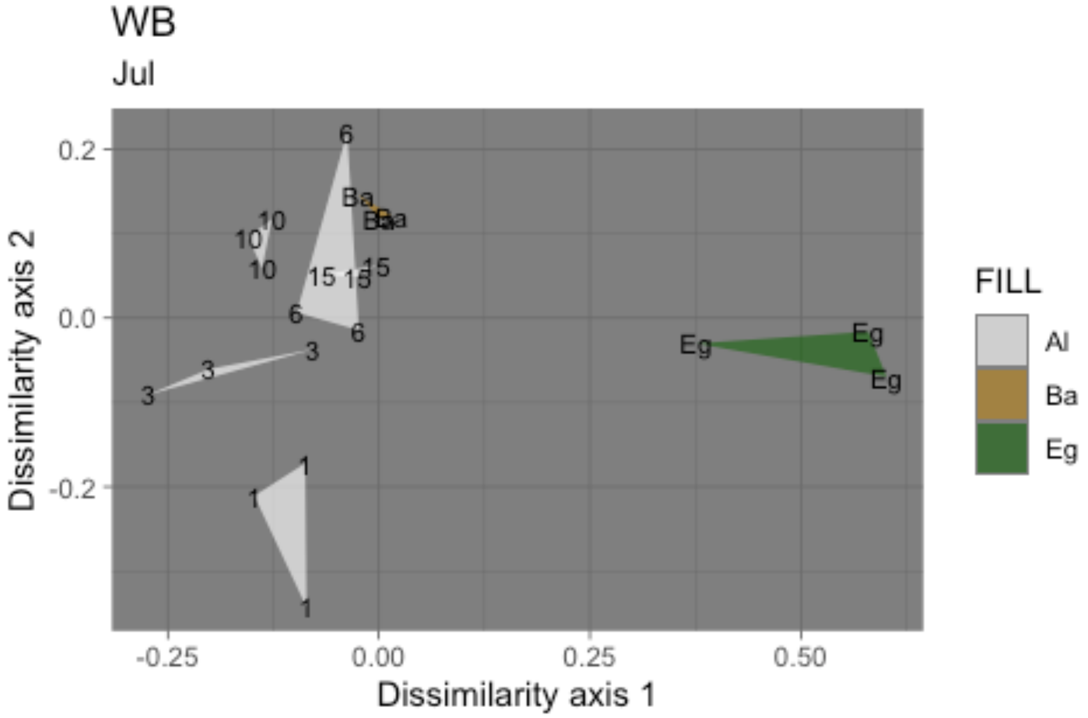


*Figure S7: Ordination plots of samples along all ten fully-sampled transects from bare to eelgrass positions. Technical replicates of each biological sample are grouped as triangles. Alongshore transect samples are shown in white and labeled with distance from the contiguous eelgrass bed in meters (Al); the single sample taken above eelgrass (located 47-90 meters inside the edge of the contiguous beds) is shown in green (Eg), and that taken above bare substrate (located 16-670m outside the edge of the contiguous beds) is shown in brown (Ba).*

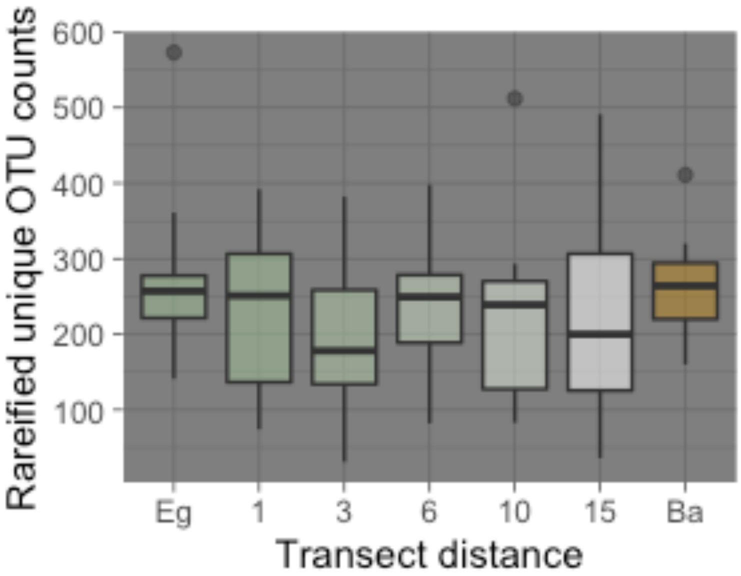


*Figure S9: OTU Richness plotted for all sites and months at each point along the transect from eelgrass to bare substrate. Bars shown are means, boxes depict the two inner quartiles, whiskers extend to the largest value no further than 1.5 times the inter-quartile range, and data outside this range are plotted individually.*
